# A unique spliced varicella-zoster virus latency transcript represses expression of the viral transactivator gene 61

**DOI:** 10.1101/174797

**Authors:** Daniel P. Depledge, Werner J. D. Ouwendijk, Tomohiko Sadaoka, Shirley E. Braspenning, Yasuko Mori, Randall J. Cohrs, Georges M. G. M. Verjans, Judith Breuer

## Abstract

During primary infection, neurotropic alphaherpesviruses (αHVs) gain access to neurons in sensory and cranial ganglia establishing lifelong latent infection from which they can later reactivate to cause debilitating disease^1^. For most αHVs, including the best-studied herpes simplex type 1 ( HSV-1), viral latency is characterized by expression of a single or restricted set of transcripts that map antisense to the open reading frame (ORF) homologous to the major HSV immediate early viral transactivator, ICP0^2^. These latency transcripts, either directly or through encoded miRNAs or proteins, repress expression of the ICP0 orthologues^3–5^. The exception is varicella-zoster virus (VZV), an αHV which infects over 90% of adults and for which neither a canonical latency transcript^1,6–8^ nor a putative mechanism for repressing lytic transcription during latency have been identified. Here, we describe the discovery and functional characterization of a VZV latency transcript (VLT), that maps antisense to VZV ORF 61 (the VZV ICP0 homologue^9,10^), and which is consistently expressed in neurons of latently infected human trigeminal ganglia (TG). VLT encodes a protein with late kinetics during lytic VZV infection *in vitro* and in zoster skin lesions. Whereas multiple alternatively spliced VLT isoforms are expressed during lytic VZV infection, a single unique VLT isoform that specifically suppresses ORF61 gene expression predominates in latently VZV-infected human TG. The discovery of VLT directly unifies the latent VZV transcription program with those of better-characterized αHVs, removing longstanding barriers to understanding VZV latency and paving the way for research into the development of vaccines that do not establish latency or reactivate, and drugs that eradicate latent VZV.

---

Primary VZV infection typically causes varicella (chickenpox), establishes a lifelong persistent infection in ganglia and reactivates in about one-third of latently infected individuals to cause herpes zoster (shingles), often accompanied by multiple serious neurological sequelae^11^. Like other herpesviruses, lytic VZV infection is characterised by full viral gene expression occurring with temporally linked immediate-early (IE), early (E) and late (L) kinetics to generate infectious virus progeny^12,13^. By contrast VZV gene expression during latency remains poorly defined^6,8,14–16^. A lack of appropriate animal^17^ and *in vitro*^18,19^ models has led to VZV latency being studied in naturally infected human TGs recovered at post-mortem. Using PCR primers targeting all canonical VZV genes, only ORF63 is occasionally detected in TGs with PMI <9 hrs^8^, whereas multiple virus transcripts of different kinetic classes are observed in human TGs with PMI >9 hrs^6,8,15^. Moreover, reports of viral protein detection by immunohistochemistry (IHC) have been largely attributed to the presence of non-VZV antibodies^16^. The lytic VZV gene ORF63, although conserved is not expressed by other latent αHVs and does not share the characteristics of other canonical αHV latency transcripts.

The aim of this study was to use RNA sequencing (RNA-seq) to map the latent transcriptome of VZV in VZV and HSV-1 co-infected human TG harvested with short PMI (≤8 hrs). Because both viruses are estimated to establish latency in <1-5% of TG neurons^1^, we increased the sensitivity of our RNA-seq approach by introducing a hybridization-based target-enrichment step for viral nucleic acids using custom overlapping 120-nucleotide RNA probes^20,21^. Co-enriching for HSV-1 transcripts allowed us to benchmark our findings against those for HSV-1 latency which has been extensively studied^1^. Our experimental approach was validated using lytically VZV- or HSV-1-infected human retinal pigmented epithelial (ARPE-19) cells to demonstrate detection of all currently annotated VZV and HSV-1 genes at high specificity and sensitivity (Fig. 1, and Supplementary Fig. 1). RNA-seq of unenriched RNA libraries from two dually infected TGs (donors 1 and 2, Supplementary Tables 1 and 2), selected for polyadenylated transcripts or ribosomal RNA-depleted total RNA (Supplementary Fig. 2), confirmed expression of the stable 1.5/2.0-kb latency-associated transcript (LAT)-derived introns, the hallmark of HSV-1 latency^22^. Enrichment for polyadenylated HSV-1 sequences in the same TGs, and five additional dually infected TGs (donors 1-7, Supplementary Tables 1 and 2), revealed both LAT introns and the near-complete 8.3-kb full-length LAT transcript from which they derive, but no other viral transcripts (Supplementary Figs. 3 and 4). The latency-associated miRNAs (mir-H2, mir-H3, mir-H4, mir-H6, mir-H7 and mir-H14)^5^ were also detected in three TGs (donors 1, 3 and 8) selected for and analysed by miRNA sequencing of unenriched RNA libraries (Supplementary Fig. 5). These data illustrate the high specificity and sensitivity of our target-enriched RNA-seq methodology and clearly demonstrate that the HSV-1 latency transcriptome in human TG is limited to the LATs and encoded miRNAs.

**Figure 1.**
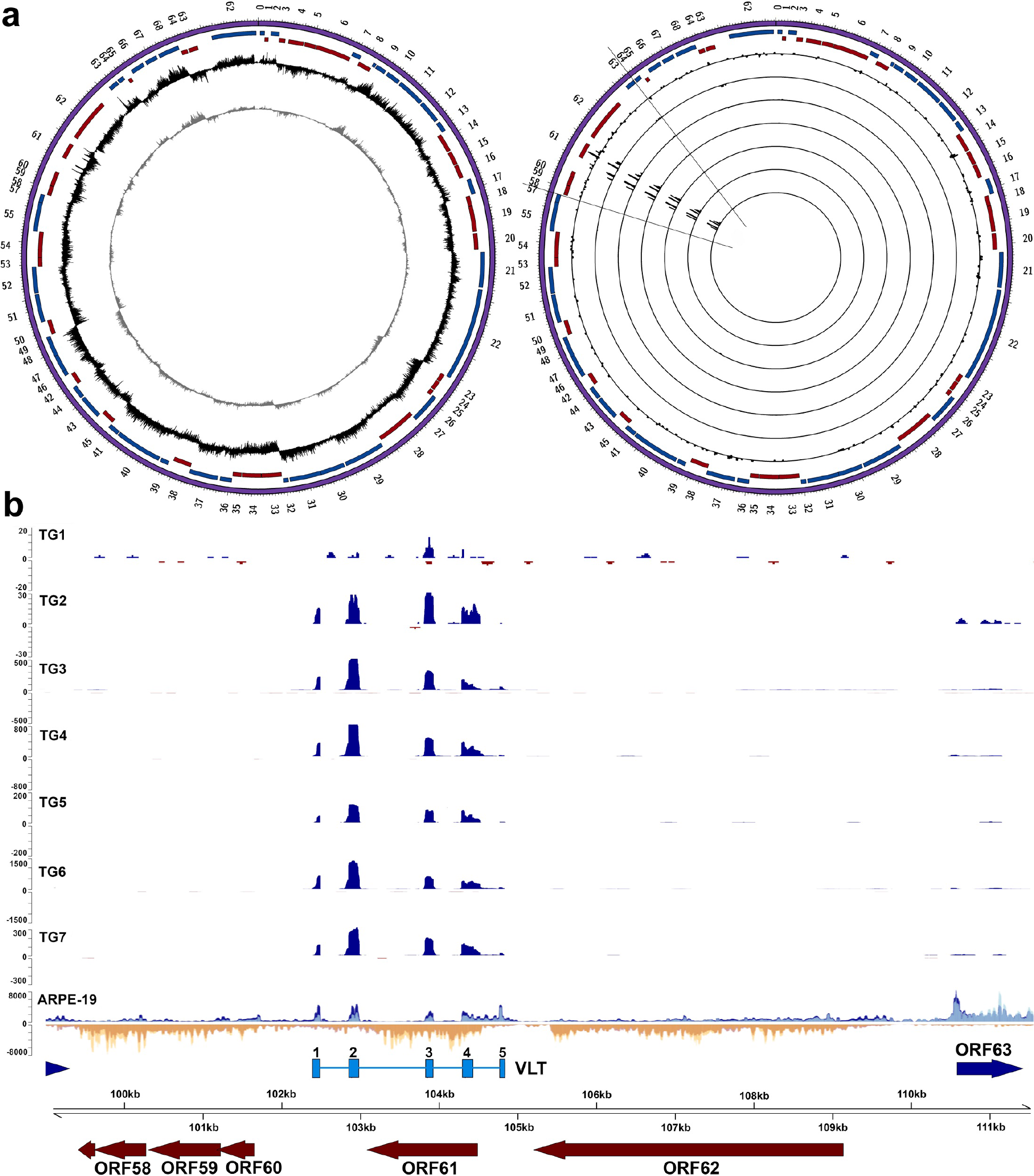
VZV transcriptome profile during lytic and latent infection. Stranded mRNA-seq of lytically VZV-infected ARPE-19 cells and seven latently VZV-infected human trigeminalganglia (TG) (Supplementary Table 1). **a**, Circos plots of the VZV genome [purple band; sense and antisense open reading frames (ORFs) indicated as red and blue blocks, respectively], with internal tracks revealing the lytic (left) and latent (right) transcriptomes using unenriched (grey track, left panel) and VZV-enriched (black tracks, left and right panels) libraries. Right panel: latent VZV transcriptome of seven TG, with each track depicting a single specimen. Peaks facing outward from the center indicate reads mapping to the sense strand, while peaks facing inward originate from the antisense strand. The y-axis is scaled to maximum read depth per library in all cases. **b**, Linear representation of the varicella latency transcript (VLT) genomic region (black lines in **a**), with blue and yellow tracks depicting VZV-enriched mRNA-Seq reads originating from the sense and antisense strand, respectively. Unenriched mRNA-Seq tracks for ARPE-19 cells, and TGs 1 and 2, are superimposed and shown in light blue (sense) or yellow (antisense), with overlapping regions in medium-blue and orange, respectively. No VZV-mapping reads were obtained from unenriched sequence datasets generated from TGs 1 and 2. VZV genome coordinates are shown the VZV reference strain Dumas (NC_001348.1); blue and red arrows indicate previously described VZV ORFs, and light blue boxes indicate the five VLT exons. Paired-end read datasets were generated with read lengths of 2х34 bp (ARPE-19) or 2х76 bp (TG1 and TG2) or 2х151 bp (TG3 - TG7).

While no VZV transcripts could be identified in non-enriched TG RNA samples (Supplementary Fig. 6), enrichment for VZV sequences in polyadenylated RNA revealed the presence of a novel 496-nucleotide multi-exon transcript expressed partially antisense to VZV ORF61 in all seven TGs analysed (donors 1-7; Fig. 1, Supplementary Figs. 6 and 7, Supplementary Tables 1 and 2). Manual inspection of the VZV sequence read data, combined with *de novo* transcript reconstruction, revealed five distinct exons (Fig. 1 and Supplementary Fig. 8a), two of which encode cleavage factor I-binding motifs (TGTA), while the most 3’ exon contained a canonical polyadenylation signal site (AAUAAA) (Supplementary Fig. 8b). We term this novel spliced transcript the VZV latency transcript (VLT). Except for ORF63 transcripts detected in six of seven TGs (donors 2-7), no other VZV transcripts (Fig. 1b and Supplementary Fig. 7) or miRNAs were detected in human TGs (data not shown). VLT was also transcribed in lytically VZV-infected ARPE-19 cells (Fig. 1b and Supplementary Fig. 7), enabling rapid amplification of cDNA ends (RACE) to confirm 5’ and 3’ transcript boundaries. Unlike the single VLT isoform detected in latently VZV-infected human TG (Fig. 2a), gene transcription from the VLT locus appears extremely complex during lytic VZV infection, with additional upstream exons or alternative splicing in exons 3 and 4 (Fig. 2b). RT-qPCR did not detect any of the three most abundant lytic VLT isoforms (each of which uses a different upstream exon) in latently VZV-infected TGs (Fig. 2c). Neither were additional upstream exons or alternative splicing of exons 3 and/or 4 evident in human TG-derived VZV RNA from re-examination of the RNA-seq data. Thus, a unique VLT isoform is selectively expressed in latently VZV-infected human TG with short PMI.

**Figure 2.**
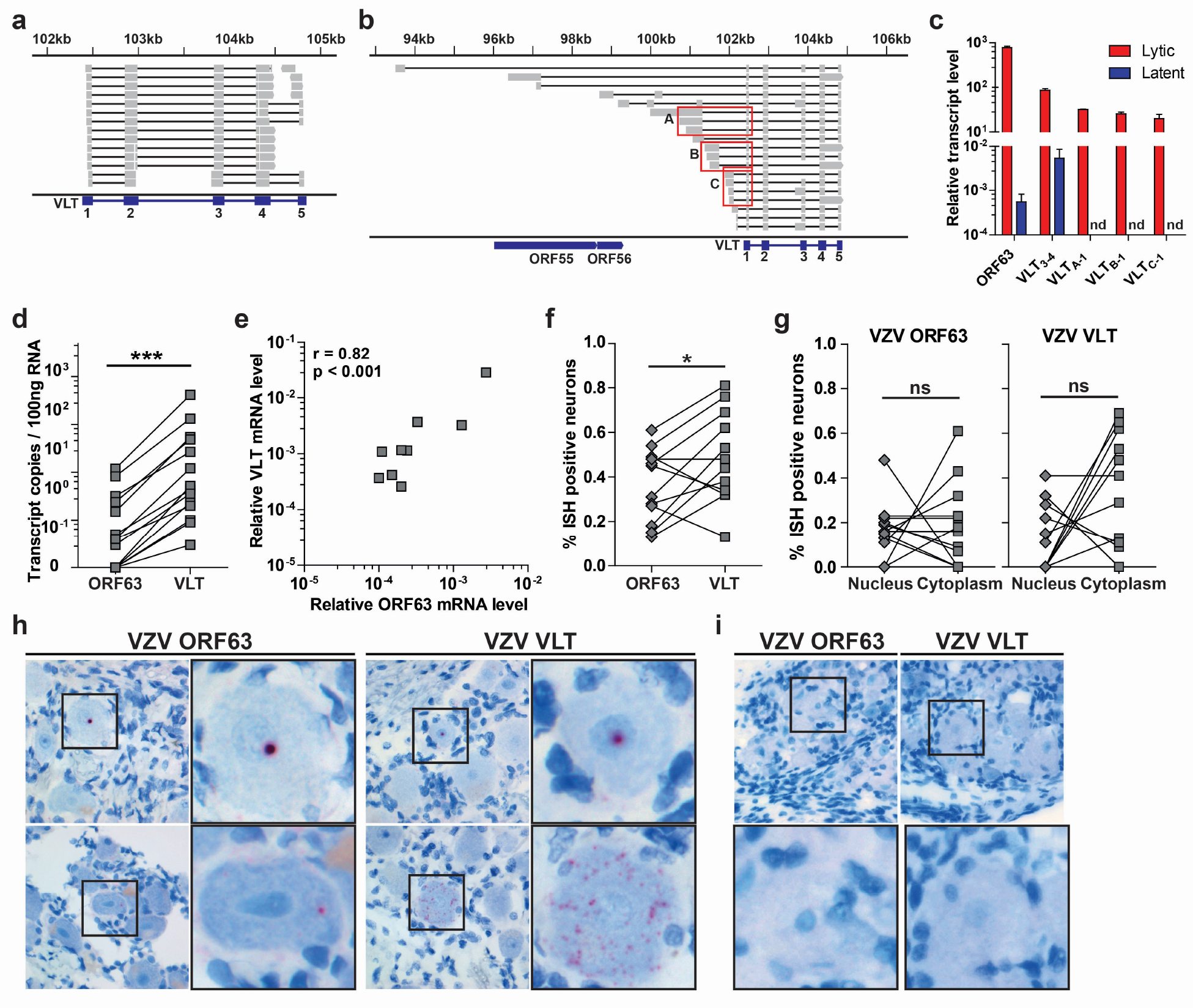
Prevalence of VZV ORF63 transcript and a unique VLT isoform in latently VZV-infected human TG. **a**, In latently VZV-infected human TG, selected paired-end reads originating from VLT sequence fragments that span multiple exons evidenced the usage of conserved splice sites between exons. Grey boxes connected by horizontal black lines indicate individual read pairs (1 – 3 per line) and show mapping regions (corresponding to the blue VLT exons), while black lines indicate skipped (spliced) regions. Data from one of four TG analysed are shown. **b**, In lytically VZV-infected MeWo cells, Sanger sequencing of amplicons generated through rapid amplification of cDNA ends (RACE) revealed multiple VLT isoforms in these cells. Grey filled boxes indicate VLT exons with splice junctions shown by horizontal black lines. Redboxes indicate most abundant groups of 5’ end splice junctions (boxes A – C), for which the percentage of recombinan*E. coli* colonies was 35% (box A), 19% (box B)and 15% (box C), as determined by colony sequencing of *E. coli* transformed with plasmids encoding PCR products spanning the 5’ end to VLT exon 5 (n=29). **c**, Detection of ORF63 and VLT isoforms in lytically VZV-infected ARPE-19 cells (red) and latently VZV-infected human TG (blue; n=19 TG) (Supplementary Table 1), using primers/probes spanning VLT exons 3⟶4, A⟶1, B⟶1 and C⟶1 (Supplementary Table 5). Data represent mean (± standard error mean; SEM) relative transcript levels normalized to β-actin RNA abundance. nd, not detected. **d**, Levels of paired ORF63 transcript and VLT (primers/probe spanning VLT exons 2⟶3) in the same VZV nucleic acid positive human TG (VZV^POS^; n=15), as determined by reverse transcription-linked quantitative PCR (RT-qPCR). *** p< 0.001; Wilcoxon signed rank test. **e**, Correlations (Spearman) between relative ORF63 RNA and VLT levels in VZV^POS^ human TG (n=15), as determined based on RT-qPCR. **f-i**, Analysis of ORF63 RNA and VLT abundance in latently VZV-infected TG (n=12) by *in situ* hybridization (ISH). **f**, Frequency of neurons positive by ISH for ORF63 RNA and VLT in consecutive TG sections of the same donor. * p<0.05; paired Student’s t-test. **g**, Nuclear and cytoplasmic ORF63 RNA and VLT expression in consecutive TG sections from individual donors. ns, not significant. **h**, Representative images of human TG sections using probes specific for ORF63 and VLT. **i**, Absence of ORF63 RNA and VLT ISH signal in two VZV naïve human fetal dorsal root ganglia. In **h** and **i**, nuclei were stained with hematoxylin. Magnification: 400X, Inset: 400X magnification and 3X digital zoom.

To confirm that VLT expression is a general feature of VZV latency, we analysed TG specimens from 18 individuals (donors 1-18, Supplementary Table 1) for the presence of VZV and HSV-1 DNA and transcripts by qPCR and reverse transcriptase-qPCR (RT-qPCR). Thirteen TG were co-infected with VZV and HSV-1, while the remaining TG contained only VZV (donors 10 and 12) or HSV-1 (donors 16 – 18), (Supplementary Table 1). Fourteen of 15 (93%) VZV nucleic acid-positive (VZV^POS^) TGs expressed VLT and 9 of 15 (60%) VZV^POS^ TGs coexpressed ORF63 mRNA at lower levels relative to VLT (Fig. 2d, Supplementary Fig. 9a and Supplementary Table 1). VLT levels correlated significantly with ORF63 transcript levels (Fig. 2e and Supplementary Fig. 9b), but not with VZV DNA load or PMI excluding the possibility of viral reactivation after death (Supplementary Figs. 9c and d)^8^.

Next, we investigated the expression and localization of VLT, and as a control VZV ORF63, in latently VZV-infected TGs (n=12) and two VZV naïve foetal dorsal root ganglia (DRG) by *in situ* hybridization (ISH). VLT and ORF63 transcripts were localized to both the neuronal nucleus and cytoplasm of distinct neurons in latently VZV-infected TG, but not in uninfected DRG (Fig. 2f-i, Supplementary Figs. 10 and 11). More TG neurons expressed VLT (0.49 ± 0.20%; mean ± SD) than ORF63 transcript (0.36 ± 0.16%) (Fig. 2f). RNase but not DNase treatment abolished ORF63- and VLT-specific ISH staining (Supplementary Fig. 10), confirming detection of VZV transcripts and not viral genomic DNA. Although previous reports suggest that TG neurons are latently infected by VZV and HSV-1 at similar frequencies^23^, both RT-qPCR and ISH (Supplementary Table 1, Supplementary Fig. 11) revealed a much higher abundance and prevalence of HSV-1 LAT compared to VLT and VZV ORF63 RNA in human TG specimens.

*In silico* translation of the VLT isoform expressed in human TG predicted a 137-amino acid protein (pVLT), with a start codon and polyadenylation site/stop codon in exons 2 and 5, respectively (Supplementary Fig. 8b). A polyclonal pVLT-specific antibody, generated against the first 19 N-terminal residues of pVLT, confirmed pVLT expression in the nucleus and cytoplasm of VLT-transfected cells and its predominantly nuclear expression in lytically VZV-infected ARPE-19 cells (Fig. 3a and b, Supplementary Figs. 8c and 12). In lytically VZV-infected ARPE-19 cells, pVLT co-localized with nuclear-expressed ORF62 protein (IE62), but not with cytoplasmically expressed glycoprotein E (Fig. 3b, Supplementary Fig. 12). The kinetic class of VLT was determined by RT-qPCR in VZV-infected ARPE-19 cells cultured in the presence or absence of phosphonoformic acid (PFA), a broad-spectrum herpesvirus DNA polymerase inhibitor^24^. Whereas no effect on ORF61 (IE gene) and ORF29 (E gene) transcription was observed, ORF49 (leaky-L gene) transcription was markedly reduced in PFA-treated VZV-infected cells (Fig. 3c). PFA blocked VLT expression completely (Fig. 3c), demonstrating that VLT transcription follows a truly late kinetic pattern *in vitro*. Finally, shingles skin biopsies were assayed for expression of VLT and pVLT. ISH revealed VLT expression throughout the affected epidermis and dermis surrounding skin vesicles (Fig. 3d), while IHC analyses of consecutive sections for VZV ORF63 protein (IE63) and pVLT indicated co-expression in the same skin areas, but not in healthy control skin (Fig. 3e, Supplementary Fig. 13). Neither pVLT- nor IE63- specific IHC signal was detected in latently VZV-infected human TG sections (data not shown).

**Figure 3.**
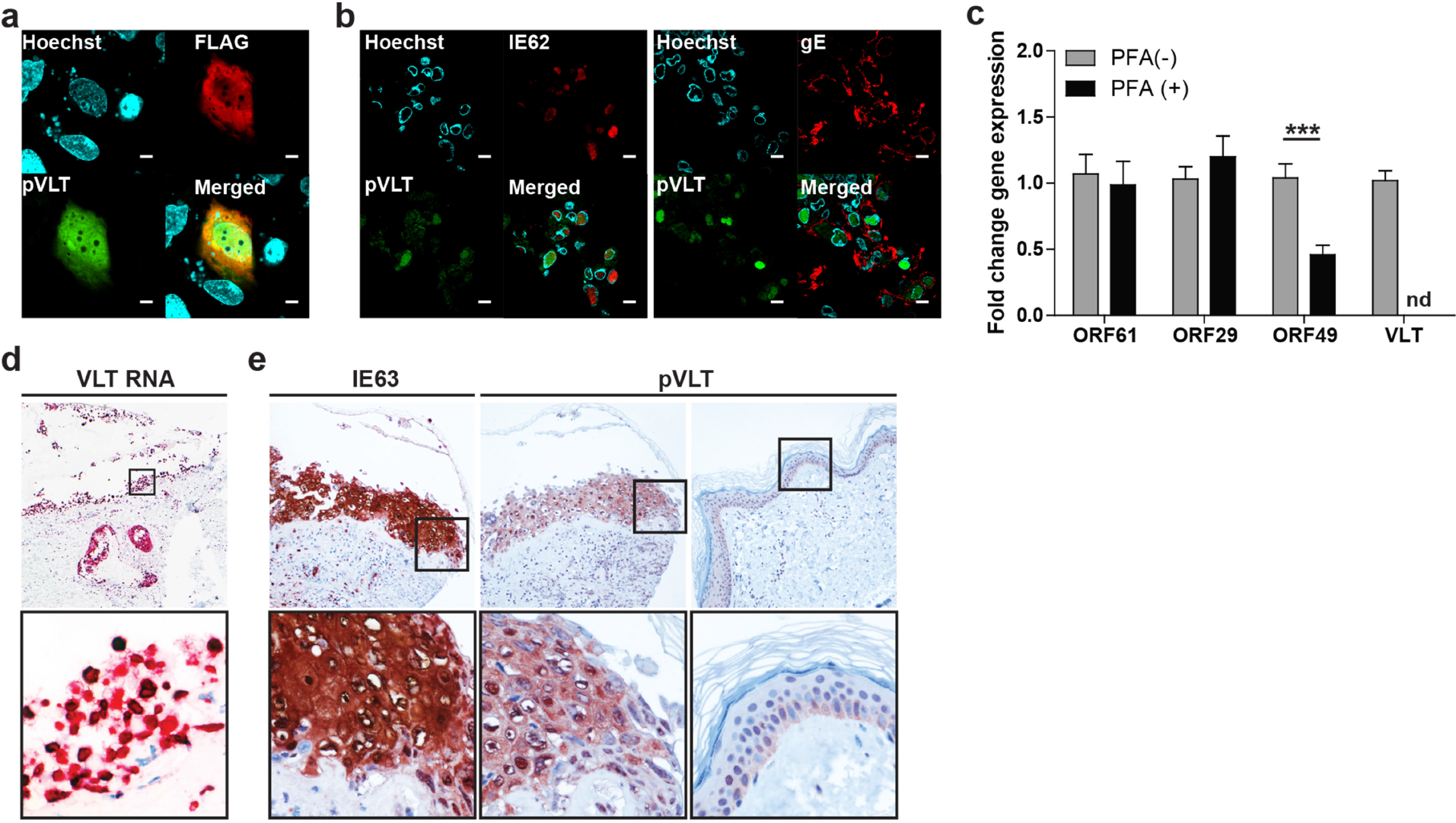
Expression and localization of VLT protein *in vitro* and *in vivo*. **a**, Representative confocal microscopy image of ARPE-19 cells at 48 hr post-transfection with C-terminal FLAG-tagged VLT (pVLT-FLAG) expression plasmid. pVLT-FLAG was detected with antibodies directed to FLAG (red) and VLT protein (pVLT; green). Magnification: 1000X and 2X digital zoom. Bars= 5 μm. **b**, Representative confocal microscopy images of VZV strain pOka-infected ARPE-19 cells at 5 days post-infection, stained for pVLT (green) and VZV ORF62 protein (IE62; red) (left panel), or pVLT (green) and VZV glycoprotein E (gE; red) (right panel). Magnification: 1000X. Bars=10 μm. In **a** and **b**, nuclei were stained with Hoechst 33342 (blue) and images are representative of results from 3 independent experiments. **c**, RT-qPCR quantitation of VZV ORF61, ORF29, ORF49 RNA and VLT levels in VZV pOka-infected ARPE-19 cells cultured with (PFA+) and without phosphonoformic acid (PFA−) for 24 hr. Data represent mean (± SEM) fold-change in gene expression, using the respective ‘PFA−’ value as a calibrator, from four independent experiments. *** p< 0.001; Wilcoxon signed rank test. **d**, Detection of VLT (red) in a human herpes zoster (HZ) skin lesion by *in situ* hybridization. Magnification: 100X. Inset: 400X and 2X digital zoom. **e**, Consecutive HZ skin sections stained immunohistochemically for ORF63 protein (IE63; brown) and pVLT (brown) (left and middle panels) and sections from unaffected control skin stained for pVLT (right panel). Magnification: 200X. Inset: 400X. Images in **d** and **e** are representative of five HZ skin biopsies stained.

Because VLT is partially antisense to VZV gene ORF61, a major viral transactivator and αHV ICP0 homologue^9^, we tested whether VLT repressed expression of ORF61 in VLT-transfected ARPE-19 cells. VLT expression significantly reduced ORF61, but not ORF62 and ORF63 transcript levels, in cells co-transfected with all four VZV genes (Fig. 4a). Western blot analysis confirmed that VLT diminishes IE61 but not IE62, IE63 and α-tubulin protein abundance in co-transfected cells (Fig. 4b). Mutation of the pVLT start codon from ATG to ATA (Supplementary Fig. 8b) resulted in loss of pVLT expression in co-transfected ARPE-19 cells, but did not abolish the inhibitory effect of VLT on ORF61 transcript and protein (IE61) expression in co-transfected cells (Fig. 4b and c, Supplementary Fig. 14). The data implicate VLT, but not pVLT, in the selective repression of VZV ORF61 gene expression.

**Figure 4.**
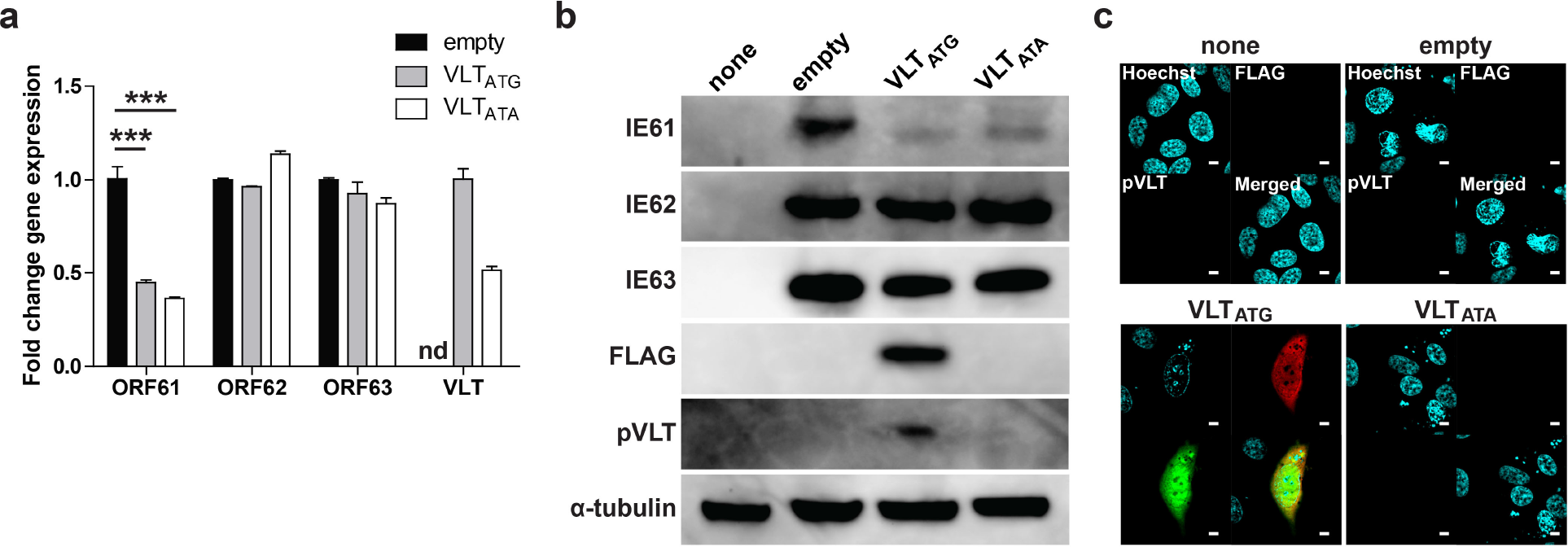
Selective repression of VZV ORF61 gene expression by VLT. ARPE-19 cells were transfected with plasmids encoding FLAG-tagged VLT (VLT_ATG_), mutated VLT in which the ATG start codon was replaced by ATA sequence (VLT_ATA_) (Supplementary Fig 8), or empty control plasmid (empty), in combination with plasmids encoding ORF61, ORF62 and ORF63. **a**, Analysis of VZV ORF61, ORF62, ORF63 and human β-actin RNA by RT-qPCR. Data represent mean (± SEM) fold-change in gene expression using the empty vector to calibrate ORF61, ORF62 and ORF63, or using VLT_ATA_ to calibrate VLT_ATG_. Data are from two independent experiments performed in duplicate. nd, not detected. *** p < 0.001; one-way ANOVA with Bonferroni’s correction for multiple comparisons. **b**, Western blot analysis using antibodies directed to proteins encoded by VZV ORF61 (IE61), ORF62 (IE62), ORF63 (IE63), VLT (pVLT) and FLAG-tagged pVLT and α-tubulin. None, untransfected ARPE-19 cells. Images are representative of 4 independent experiments. **c**, Confocal microscopy image of cells stained with antibodies directed to FLAG (red) and pVLT (green), and nuclei stained with Hoechst (blue). Magnification: 2000X. Bars=5 μm. Images are representative of two independent experiments.

In this study, we identified a novel and unique, spliced VZV transcript, VLT, which is consistently and selectively expressed in latently VZV-infected human TG neurons. The VLT locus, including splice donor/acceptor sites and pVLT coding sequence, is highly conserved among wild-type and vaccine strains of VZV (Supplementary Table 4). A feature shared between well-characterized latency transcripts of other αHVs, notably HSV-1 and bovine herpesvirus 1, is their ability to encode repressive miRNAs^5,25^. However, consistent with previous analyses of human TG^26^, we found no evidence of miRNAs encoded within VLT or the wider viral genome. In further contrast to other αHVs, latent VZV also transcribes ORF63 in a subset of latently infected TGs, suggesting a role for this gene in latency or early reactivation. We also find no evidence for expression of other VZV genes during latent infection. Moreover, unlike HSV-1 LAT, VLT encodes a protein that is expressed with true late kinetics in lytically VZV-infected cells *in vitro* and in shingles skin lesions. Whereas the function of pVLT remains unclear, we showed that VLT specifically suppresses expression of VZV IE61, a homolog of HSV-1 ICP0 and a promiscuous transactivator of lytic viral promotors^10^. Although VZV disease is now much reduced due to effective vaccines^27–29^, latent persistence of the VZV vOka vaccine strain, that like wild-type VZV can reactivate later in life, will necessitate continuing lifelong surveillance and potentially revaccination^30^. With our findings the field is now better positioned to understand the molecular pathways by which VZV establishes and maintains latency. This information will be critical for the future development of measures to enhance control of this ubiquitous human herpesvirus.

## Acknowledgements

DPD is supported by a New Investigator Award from the Medical Research Foundation (UK MRC) and a small grant provided by the Daiwa Foundation. JB was partially funded by the UCL/UCLH Biomedical Research Centre (BRC). TS received funding from the Takeda Science Foundation, Japan Society for the Promotion of Science (JSPS KAKENHI JP17K008858) and the Ministry of Education, Culture, Sports, Science and Technology (MEXT KAKENHI JP17H05816) and was, in part, supported by a Grant-in-Aid for Scientific Research on Innovative Areas from MEXT of Japan (JP16H06429 and JP16K21723). WJDO, RJC and GMGMV were partly supported by National Institutes of Health grant AG032958. GMGMV received funds from the Niedersachsen-Research Network on Neuroinfectiology (N-RENNT) of the Ministry of Science and Culture of Lower Saxony (Germany). RJC was additionally supported by Public Health Service grant NS082228. We acknowledge support from the Medical Research Council and BRC for the UCL/UCLH Pathogen Sequencing Pipeline as well as the UCL Legion High Performance Computing Facility, and associated support services, in the completion of this work. The authors would like to acknowledge Sarah Getu and Suzanne van Veen for technical assistance (Dept. of Viroscience, Erasmus MC, Rotterdam, The Netherlands) and the whole team of the Netherlands Brain Bank (www.brainbank.nl) for their work and contributions.

## Author contributions

DPD, WJDO, TS, GMGMV and JB designed the study. DPD, WJDO, TS and SEB performed experiments. All authors analysed and critiqued the data. DPD, WJDO, TS, RJC, GMGMV and JB wrote the manuscript.

## Supplementary Figures

**Supplementary Figure 1.**
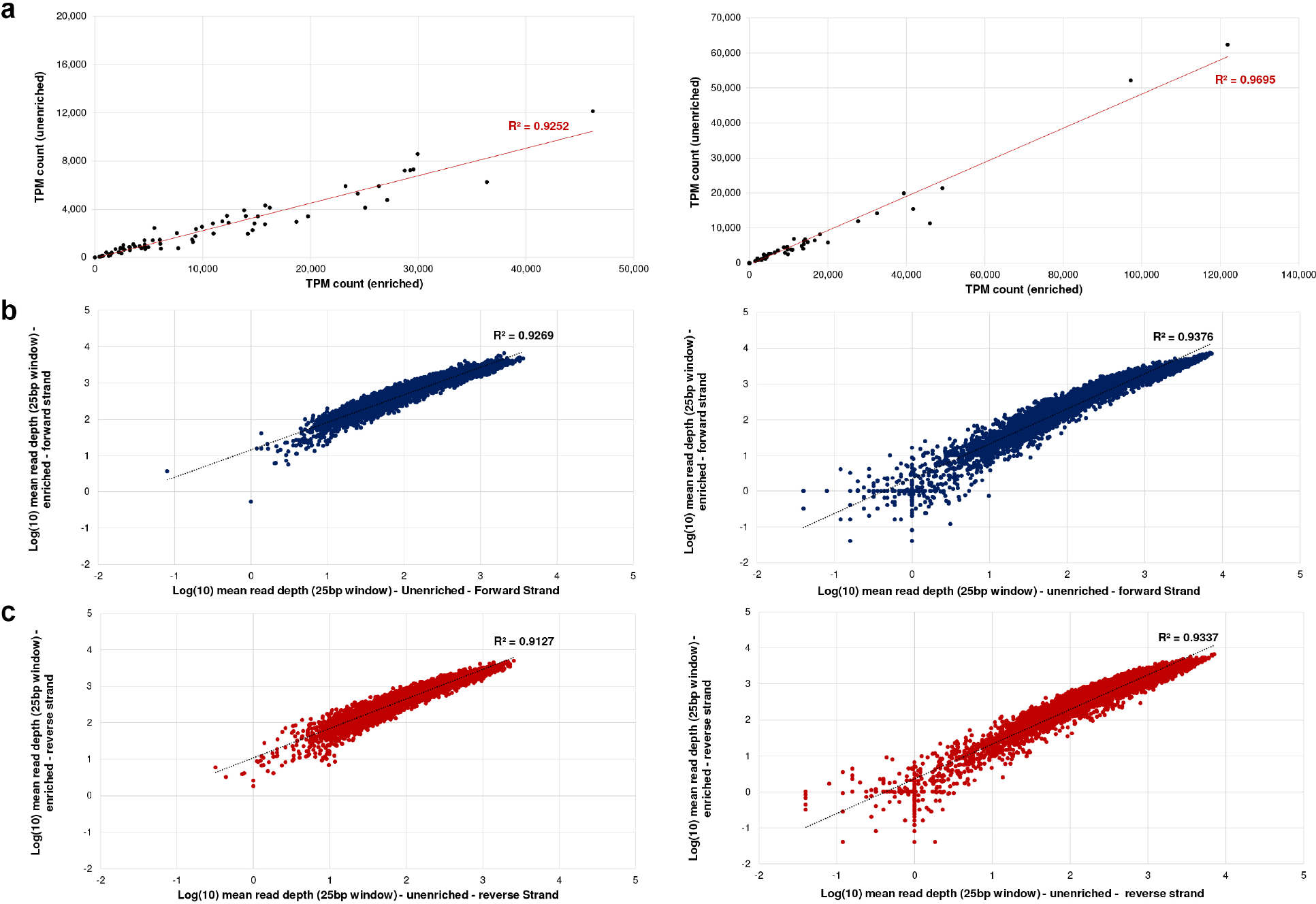
Unbiased enrichment of HSV-1 and VZV transcriptomes in lytically virus-infected human ARPE-19 cells. **a**, Transcript per million (TPM) counts were generated for all canonical ORFs of VZV (left) and HSV-1 (right) in VZV/HSV-1–enriched and unenriched sequence datasets derived from poly-A-selected RNA extracted from lytically virus-infected ARPE-19 cells. R^2^ correlation scores are shown in red. **b** and **c**, Mean read depths across iterative 25-nt windows for VZV (left) and HSV-1 (right) genomes were plotted using a log scale for VZV/HSV-1–enriched and unenriched sequence datasets derived from poly-A-selected RNA. Forward strand (**b**) and reverse strand (**c**) show a high correlation (R^2^ > 0.91) between enriched and unenriched datasets.

**Supplementary Figure 2.**
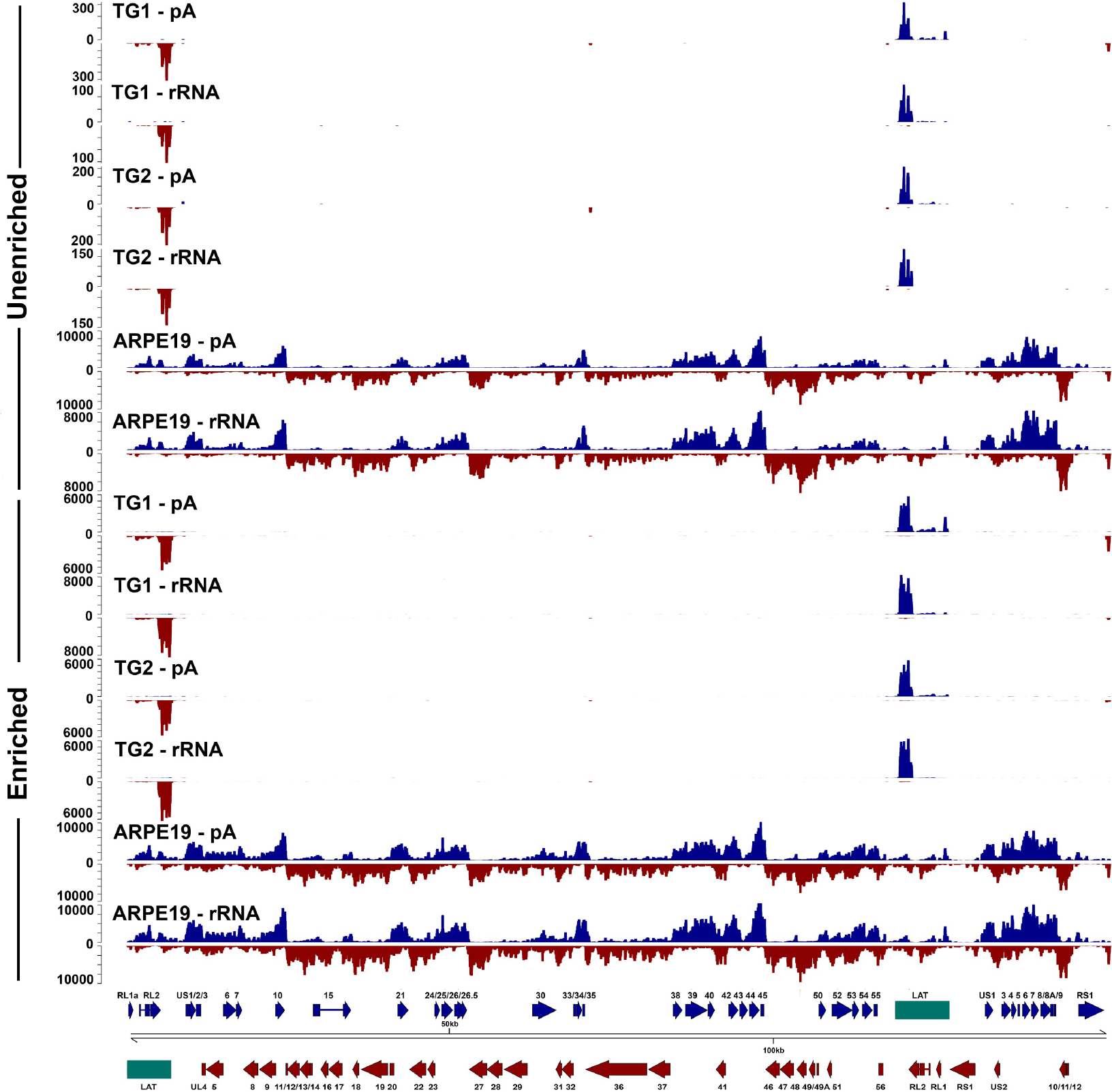
Comparison of unenriched and enriched poly-A-selected and rRNA-degraded HSV-1 transcriptome in latently HSV-1-infected human TG. RNA-Seq read data presented in a linear format along the entire 152-kb HSV-1 genome for TG specimens from donors 1 and 2 (Supplementary Table 1) and for lytically HSV-1-infected ARPE-19 cells. Each track depicts either unenriched or enriched poly-A-selected (pA) or rRNA-degraded (rRNA) HSV-1 transcriptome. Blue and red tracks represent read data originating from the sense and antisense strand, respectively. Y-axis values indicate read depth. Previously described HSV-1 ORFs within this locus (blue and red arrows) are shown. Paired-end read datasets were generated with read lengths of 2×34 bp (ARPE-19 cells) or 2×76 bp (TG1 and TG2).

**Supplementary Figure 3.**
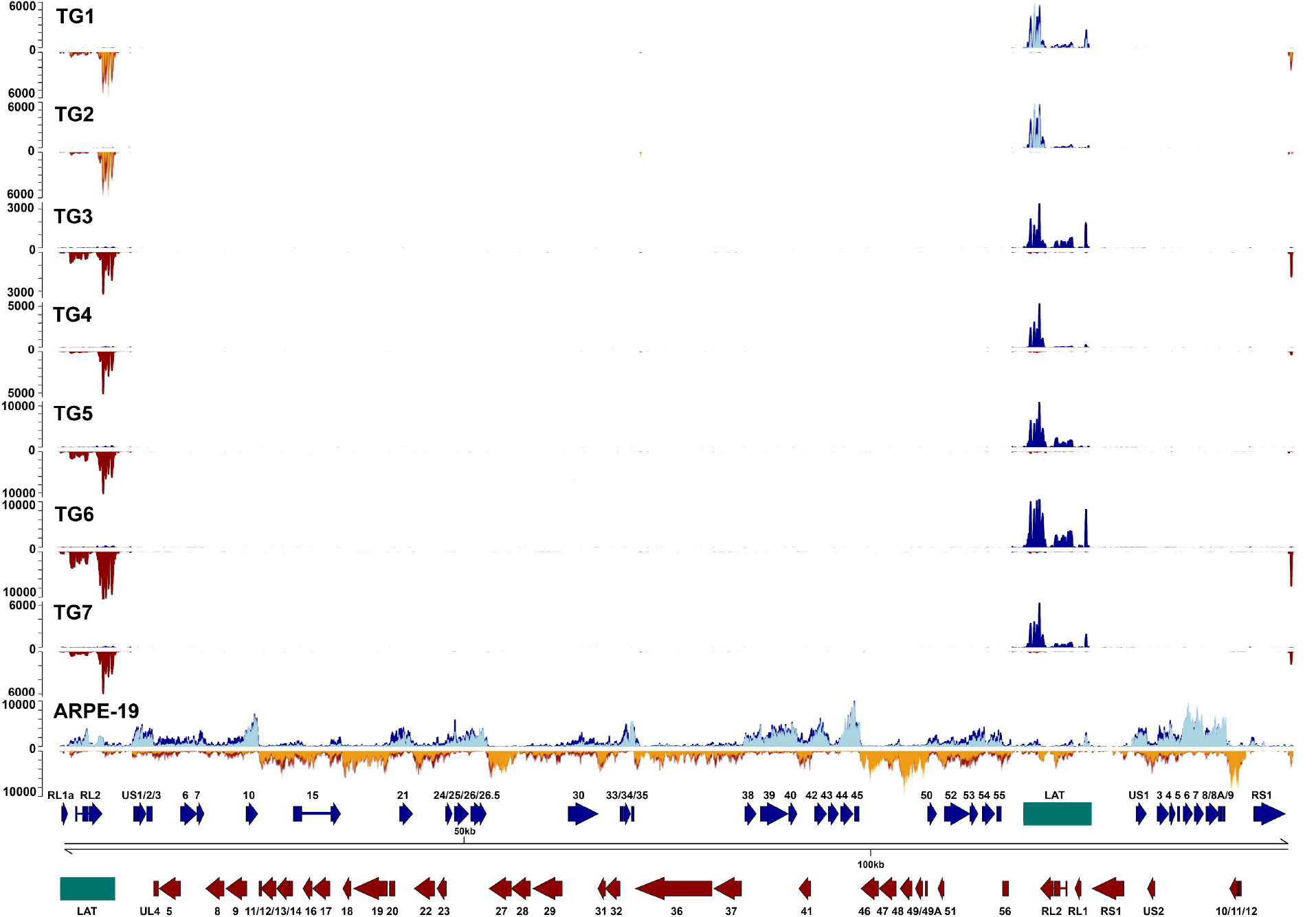
The latent and lytic transcriptome of HSV-1. RNA-Seq read data presented in a linear format along the entire 152-kb HSV-1 genome for all seven TG specimens separately and lytically HSV-1-infected ARPE-19 cells. Each track depicts the poly-A-selected, HSV-1 enriched transcriptome. Blue and red tracks represent read data originating from the sense and antisense strand, respectively. Y-axis values indicate read depth (!!label figure itself!!). Unenriched tracks superimposed on TG1, TG2 and ARPE-19 are shown in light blue (sense) or orange (antisense). Previously described HSV-1 ORFs are indicated by blue (sense strand) and red (antisense strand) arrows, while the HSV-1 latency-associated transcripts (LATs) are indicated by green boxes. Paired-end read datasets were generated with read lengths of 2х34 bp (ARPE-19), 2×76 bp (TG1 and TG2) or 2×151 bp (TG3 - TG7).

**Supplementary Figure 4.**
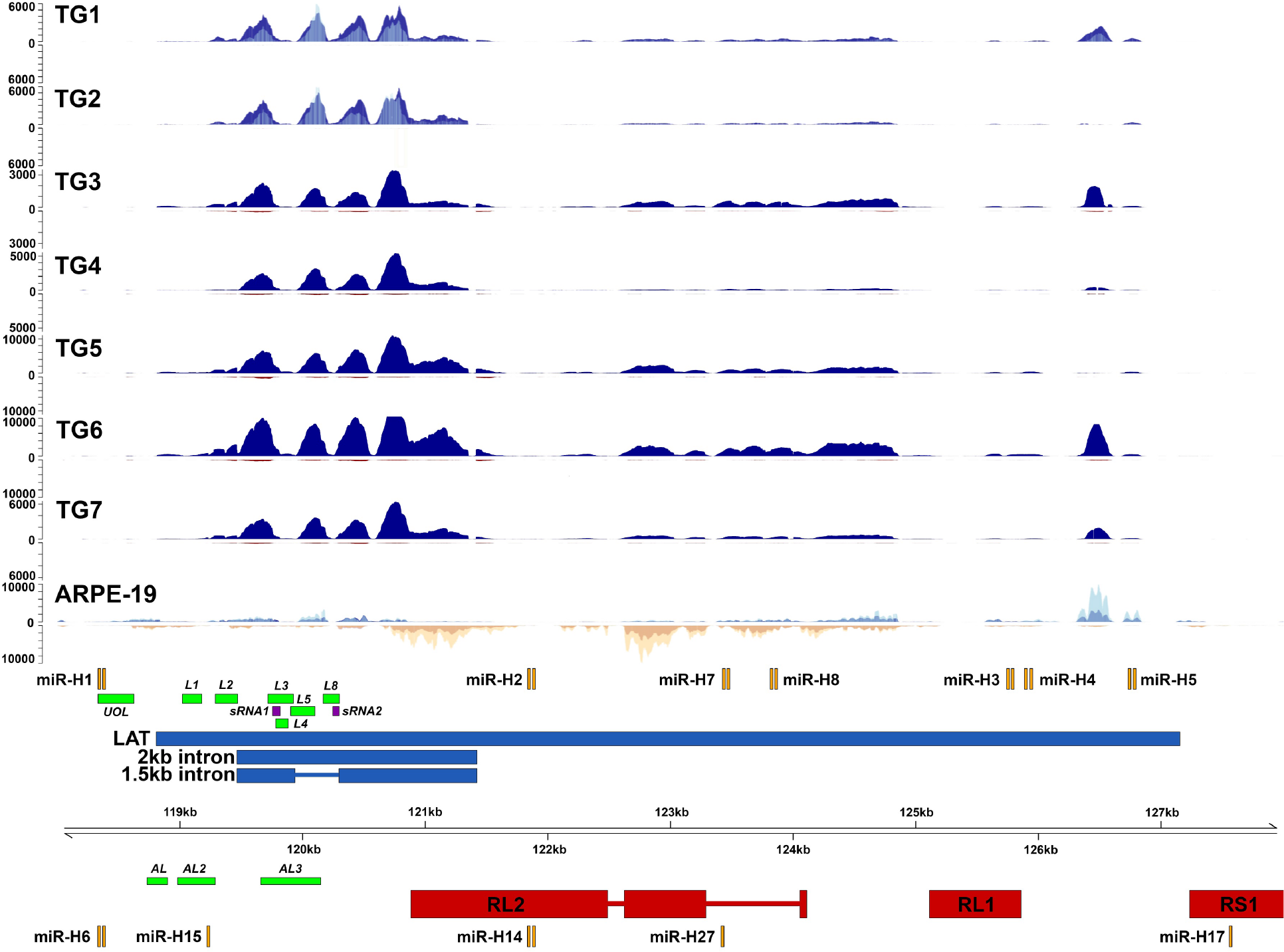
Lytic and latent transcriptome across the HSV-1 LAT locus. RNA-Seq read data presented in a linear format across the HSV-1 LAT region (sense strand coordinates 118,000-128,000 bp of HSV-1 reference strain 17; NC_001806) for each of the HSV-1-enriched transcriptomes of all seven TG specimens separately and for lytically HSV-1-infected ARPE-19 cells. Each track depicts poly-A-selected, HSV-1 enriched transcriptomes. Blue and yellow tracks represent read data originating from the sense and antisense strand, respectively. Y-axis values indicate read depth. Unenriched tracks superimposed on TG1, TG2 and ARPE-19 are shown in light blue (sense) or orange (antisense). Previously described ORFs within this HSV-1 locus (red blocks), miRNAs (orange blocks), predicted LAT-encoded ORFs (green blocks^2^), LAT-encoded small RNAs (purple blocks) and LAT (blue boxes) are scaled representatively.

**Supplementary Figure 5.**
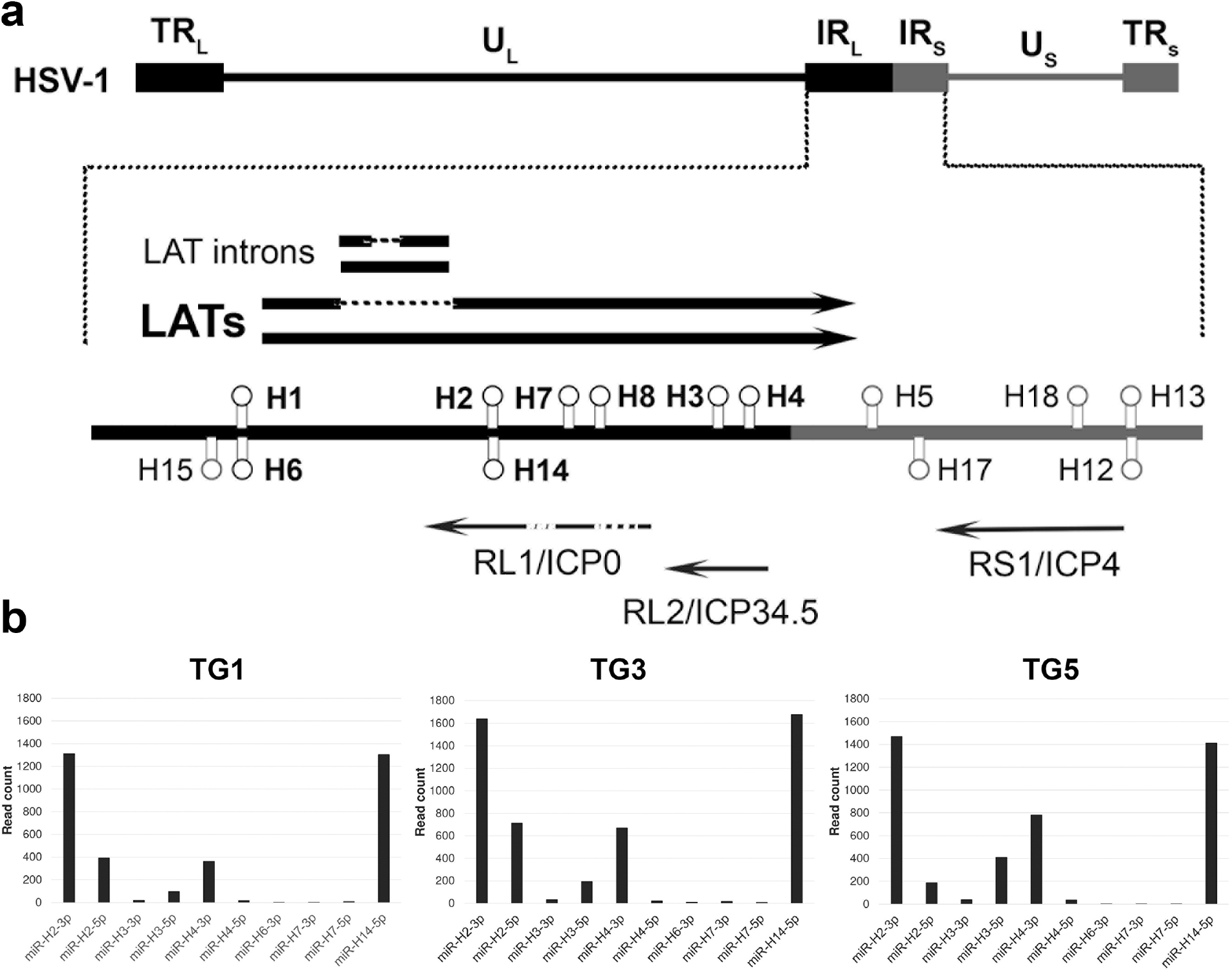
HSV-1 miRNAs detected in latently HSV-1-infected human TG. **a**, Five canonical miRNAs (miR-H2, 3, 4, 7 and 8) are encoded within HSV-1 LAT, while a sixth (miR-H14) is encoded antisense to miR-H7. Two additional miRNAs (miR-H1 and H6) expressed during latency are encoded just upstream of the 5’ end of LAT. **b**, miRNA-Seq of TG1, TG3 and TG5 confirmed transcription of these and the absence of remaining miRNAs encoded in the wider HSV-1 LAT locus (and wider genome, data not shown). Only miRNAs producing >5 reads are shown.

**Supplementary Figure 6.**
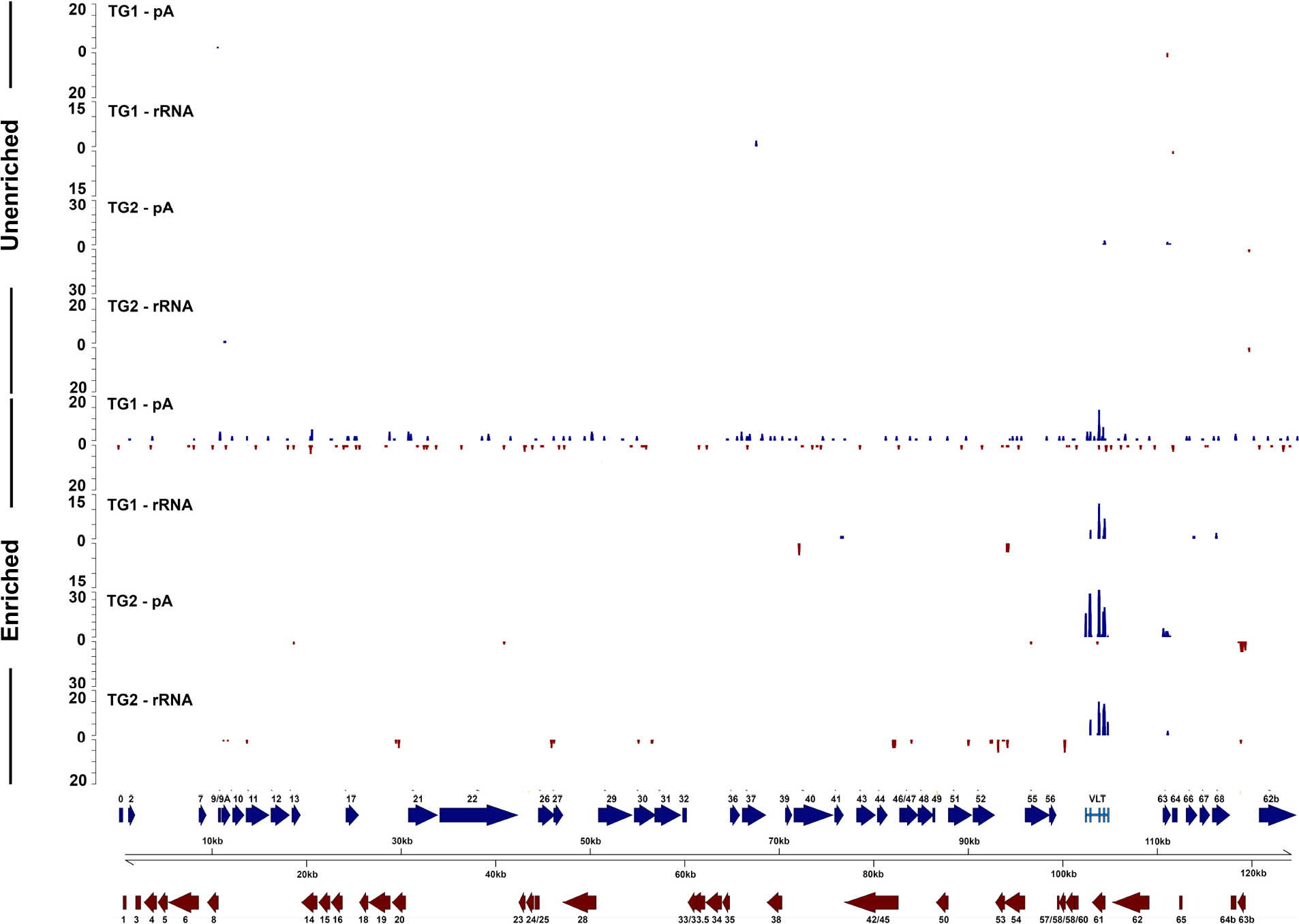
Comparison of unenriched and enriched poly-A-selected and rRNA-degraded VZV transcriptome in latently VZV-infected human TG. RNA-Seq read data presented in a linear format along the entire 125-kb VZV genome for TG specimens from donors 1 and 2 (Supplementary Table 1). Each track depicts either an unenriched or enriched poly-A-selected (pA) or rRNA-degraded (rRNA) VZV transcriptome. Blue and red tracks represent read data originating from the sense and antisense strand, respectively. *y*-axis values indicate read depth]. Previously described VZV ORFs (blue and red arrows) and the five VLT exons (light blue boxes) were scaled representatively. Paired-end read datasets were generated with read lengths of 2×76 bp.

**Supplementary Figure 7.**
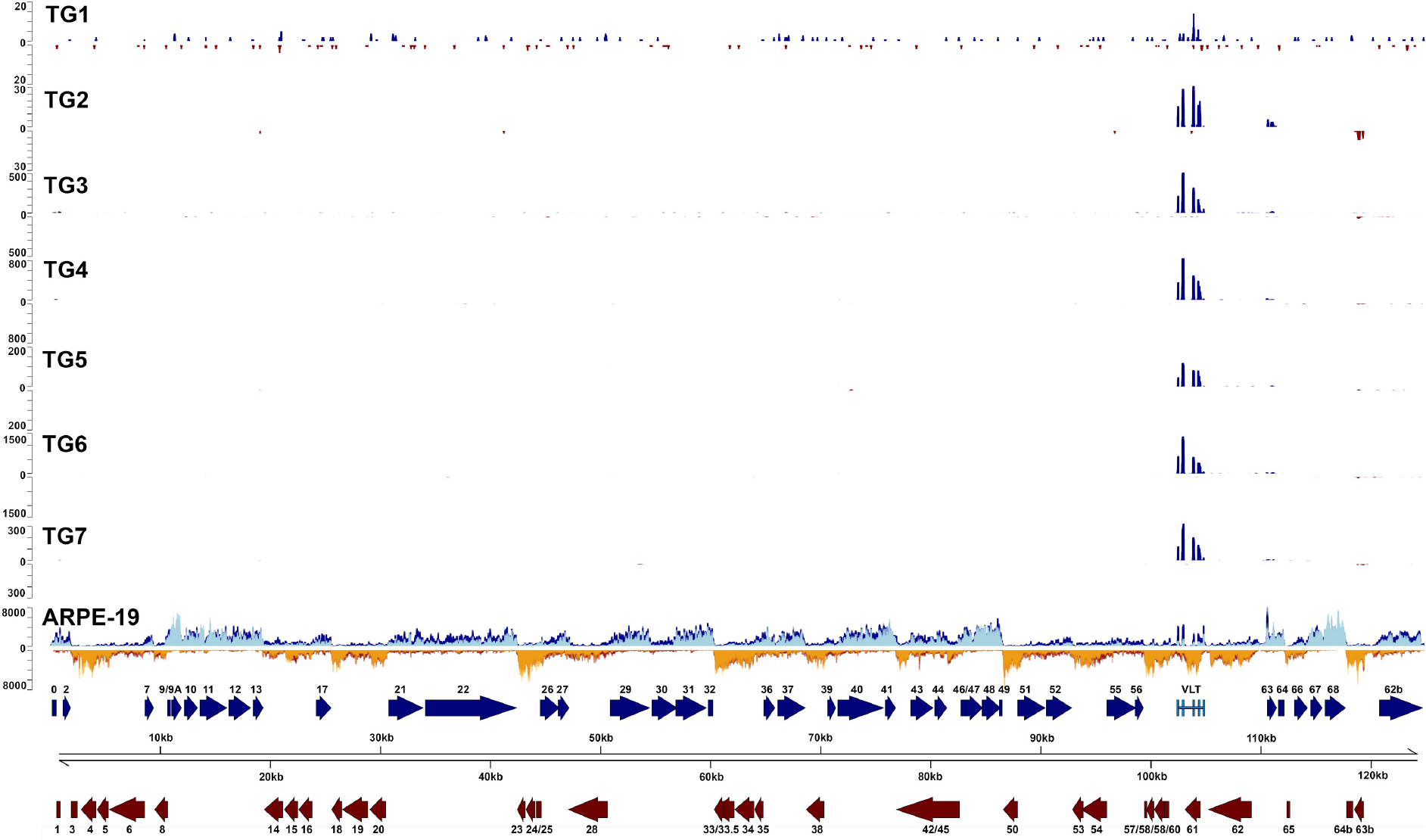
The latent and lytic transcriptome of VZV. RNA-Seq read data for seven TG specimens and lytically VZV-infected ARPE-19 cells (Fig. 1), presented in a linear format along the entire 125-kb VZV genome. Each track depicts poly-A-selected, VZV-enriched transcriptomes. Blue and red tracks represent read data originating from the sense and antisense strand, respectively. Y-axis values indicate read depth. Unenriched tracks superimposed on ARPE-19 cells are shown in light blue (sense) or orange (antisense). Note that no VZV-mapping reads were obtained from unenriched sequence datasets generated for TG1 and TG2. Previously described VZV ORFs within this locus (blue and red arrows) and the five VLT exons (light blue boxes) are scaled representatively. Paired-end read datasets were generated with read lengths of 2×34 bp (ARPE-19) or 2×7 6bp (TG1 & TG2) or 2×151 bp (TG3 - TG7). VLT was consistently detected in all TGs. Background noise observed in TG1 likely reflects a combination of low sequencing depth and low VZV genome abundance (Supplementary Table 1).

**Supplementary Figure 8.**
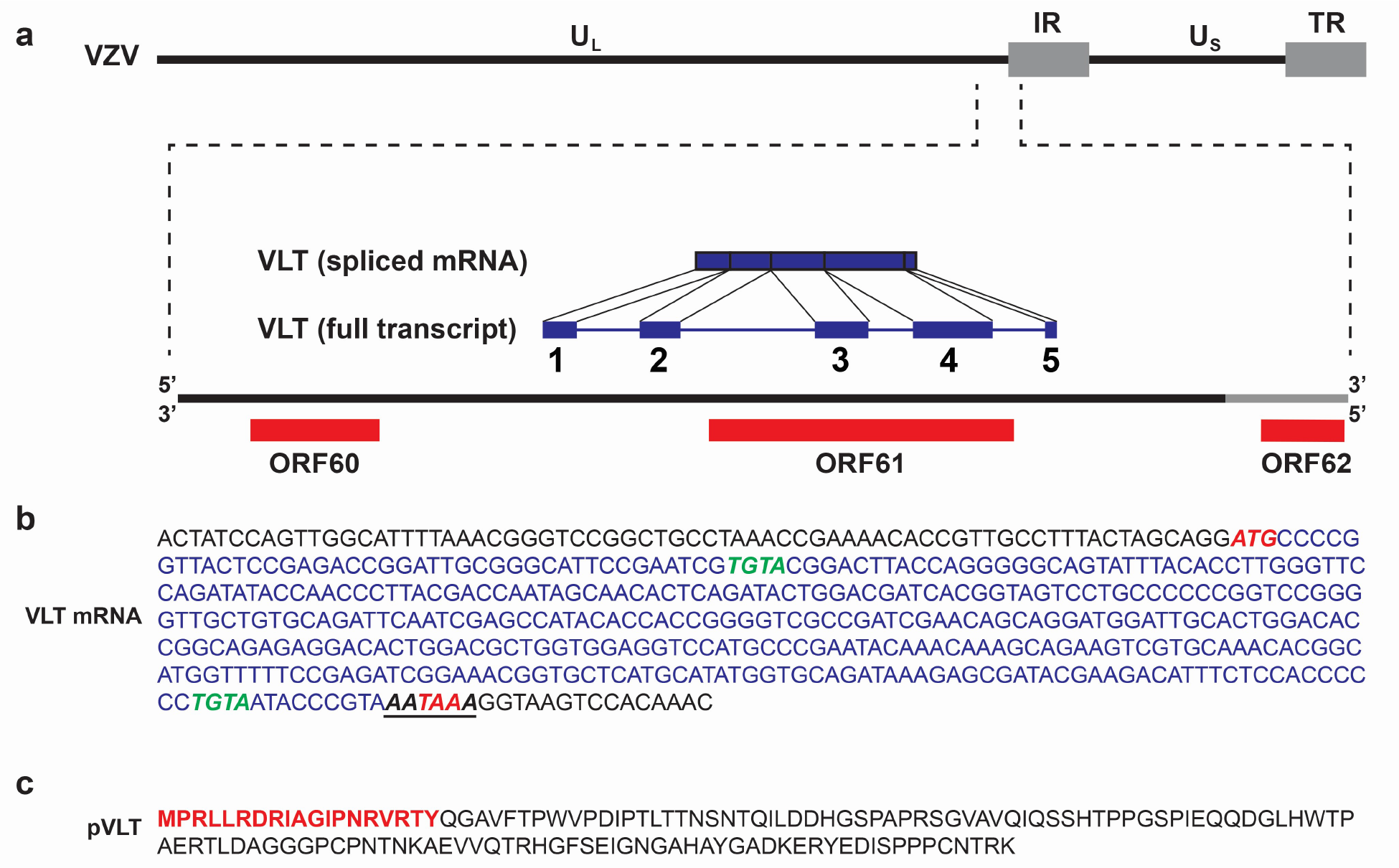
The genomic locus encoding the VZV latency transcript. **a**, Schematic diagram showing the location and structure of the five VZV latency transcript (VLT) exons (blue blocks) and introns (black lines) within the genomic region 101,000-106,000 (coordinates refer to VZV reference strain Dumas; NC_001348.1) (Supplementary Table 3). **b**, The VLT mRNA sequence including the 5’ untranslated region; start and stop codons are highlighted in red italic, while the cleavage factor I (CFI)-binding motifs are highlighted in green italic and the canonical polyadenylation signal site (AATAAA) is underlined. **c**, The fully translated VLT protein (pVLT), with the sequence of peptide used to produce rabbit polyclonal anti-pVLT antibody indicated in red.

**Supplementary Figure 9.**
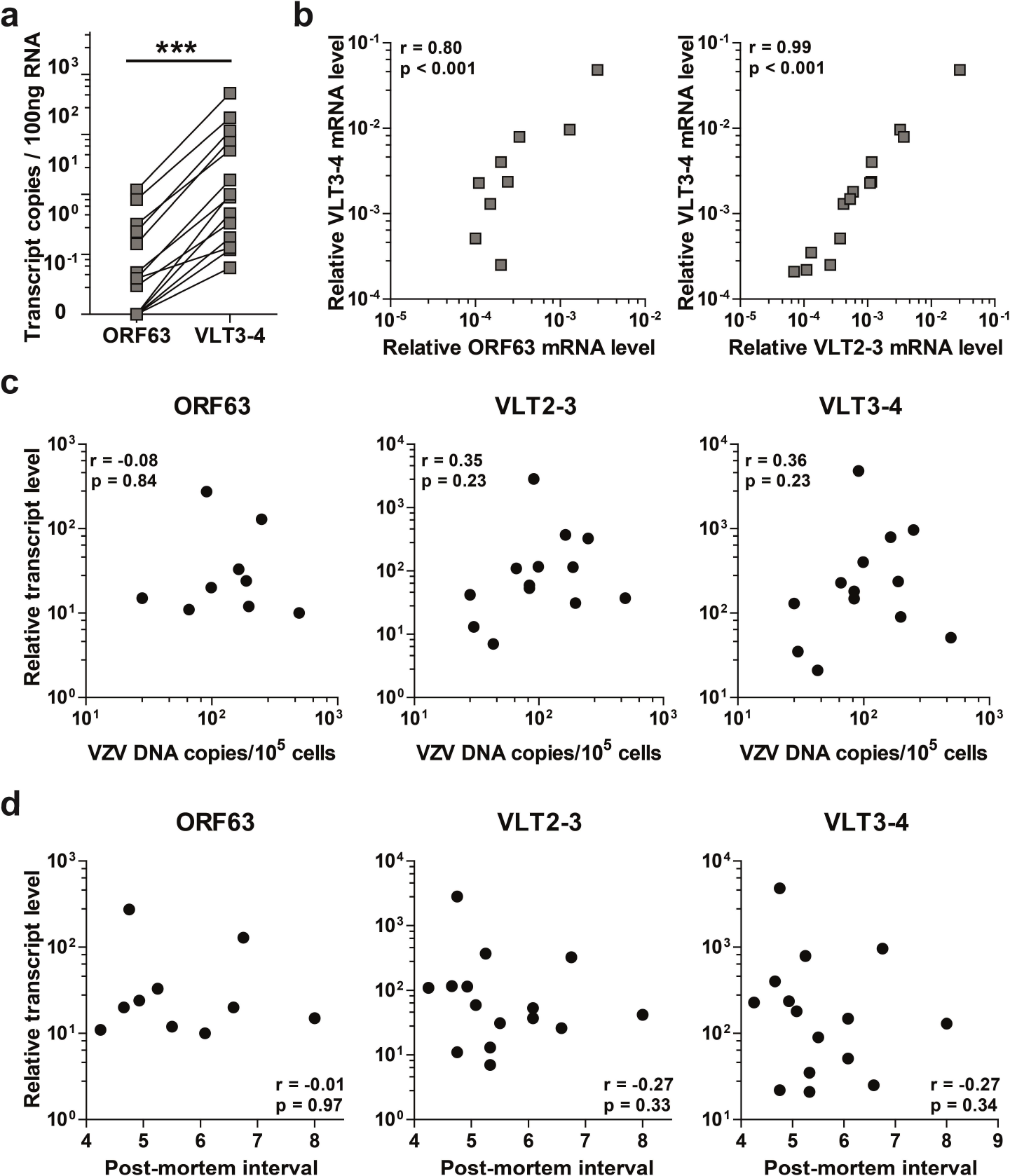
Prevalence of VZV ORF63 RNA and VLT in human TG. **a**, Relative abundance of ORF63 RNA and VLT levels in 15 VZV nucleic acid positive (VZV^POS^) human TG determined by RT-qPCR. VLT2-3 and VLT3-4 refer to primers/probes spanning the VLT exons 2⟶3 and 3⟶4 junctions (Supplementary Table 5). ORF63 RNA and VLT levels were paired in the same TG specimens. *** p< 0.001; Wilcoxon signed rank test. **b**, Correlation between relative ORF63 RNA and VLT levels, both normalized to cellular β-actin transcript levels. Spearman correlations are indicated. **c** and **d**, Correlation between relative ORF63 and VLT levels and VZV DNA load (**c**) and post-mortem interval (**d**) by qPCR in 15 VZV^POS^ human TG. Spearman correlations are indicated.

**Supplementary Figure 10.**
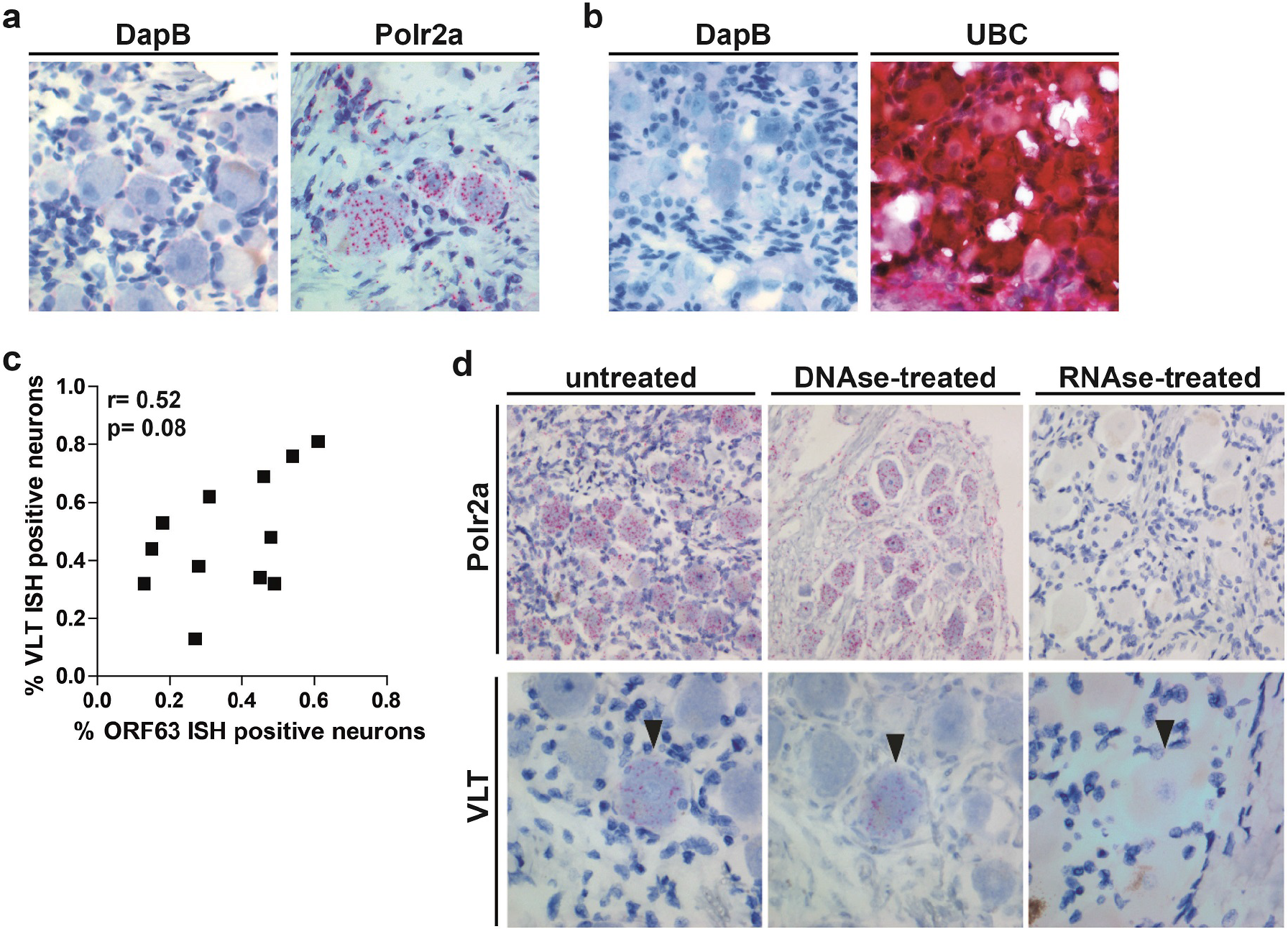
VLT-specific probe detects viral RNA, but not viral DNA in human TG by *in situ* hybridization. **a** and **b**, Analysis of negative control (bacterial gene dihydrodipicolinate reductase, DapB) and positive control [human gene RNA polymerase II polypeptide A (Polr2a) and ubiquitin C (UBC)] transcripts in human (**a**) latently VZV-infected adult TG and (**b**) VZV naïve fetal dorsal root ganglia by ISH. Magnification: 400X. **c**, Correlation between the percentages of ORF63 RNA- and VLT-positive neurons in latently VZV-infected human TG, determined by in situ hybridization (ISH). Spearman correlation is indicated. **d**, Analysis of ISH probe specificity using untreated, DNase-treated or RNase-treated consecutive sections of human TG. Magnification: 200X (Polr2a) and 400X (VLT). Arrowheads indicate the same neuron in adjacent TG sections. Representative images from two donors (n=2 sections/donor) are shown.

**Supplementary Figure 11.**
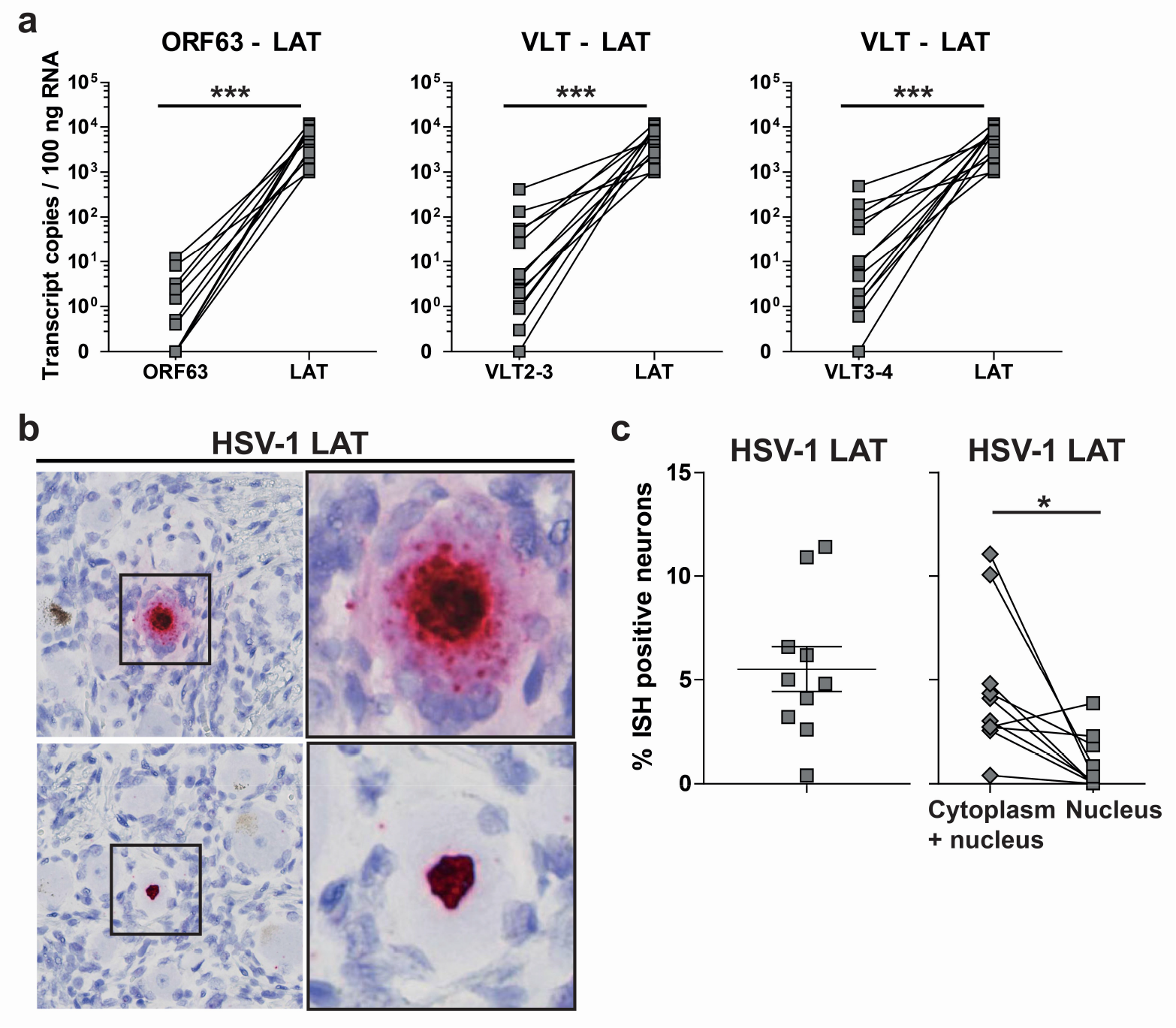
Comparison of VZV transcripts and HSV-1 LAT in human TG. **a**, qPCR analysis of HSV-1 LAT, VZV ORF63 RNA and VLT levels in 13 latently HSV-1/VZV coinfected human TG (Supplementary Table 1). VLT2-3 and VLT3-4 indicate primer/probe sets spanning VLT exons 2⟶3 and 3⟶4 junctions (Supplementary Table 5). *** p < 0.001; Wilcoxon signed rank test. **b** and **c**, Analysis of HSV-1 LAT in ten latently HSV-1-infected human TG by ISH. **b**, Representative images of ISH analysis on human TG sections using probes specific for HSV-1 LAT, showing neuronal LAT staining in both the nucleus and cytoplasm (top panels) or solely nuclear (bottom panels). Magnification: 400X; inset: 400X and 3X digital zoom. **c**, Frequency of neurons expressing HSV-1 LAT (left panel), and frequency of neurons expressing LAT in both the nucleus and cytoplasm or only in the nucleus. Horizontal line and error bars indicate average ± SEM. * p<0.05; paired Student’s t-test.

**Supplementary Figure 12.**
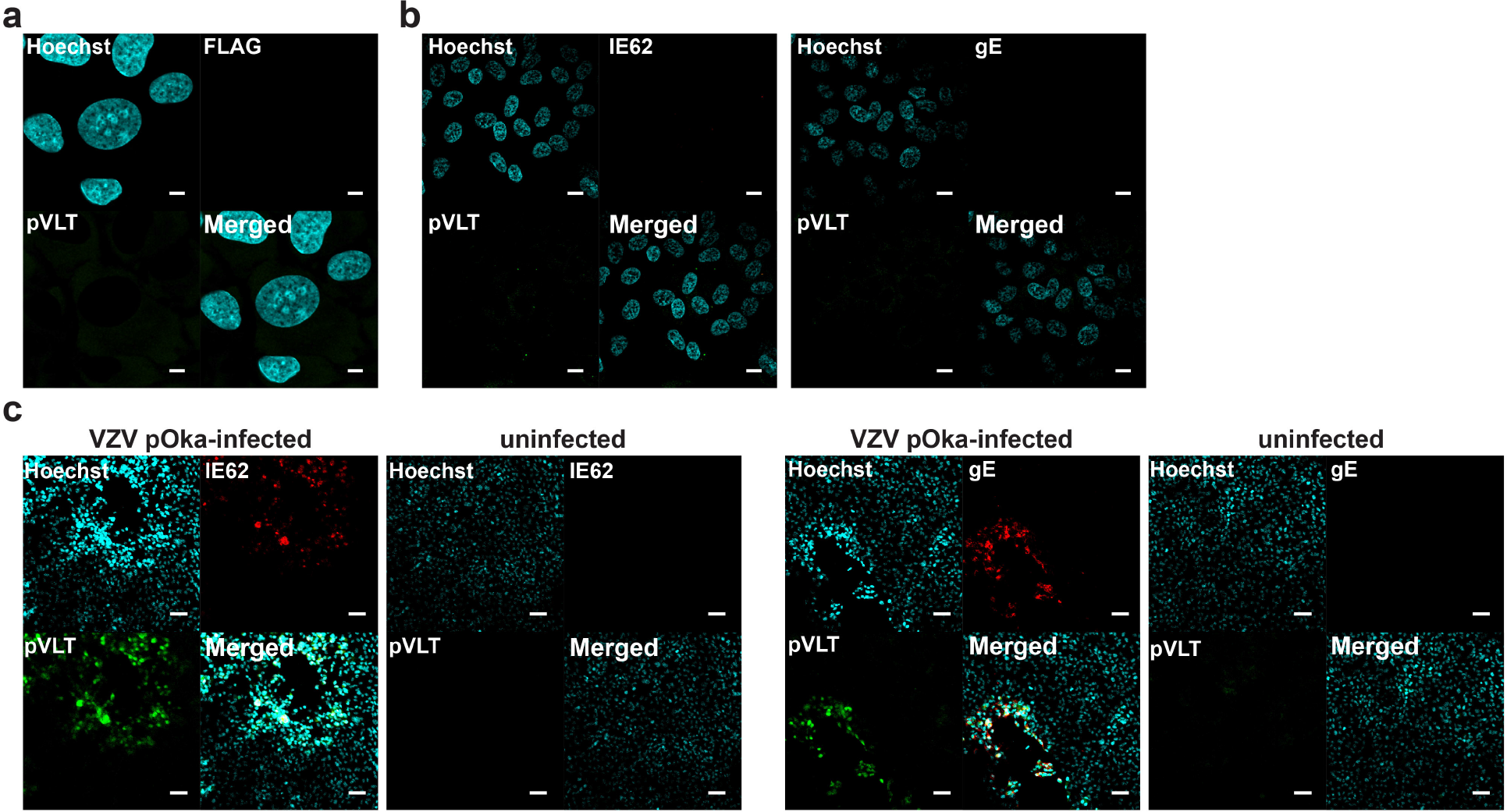
Expression of VLT protein in transfected and lytically VZV-infected cells *in vitro*. **a**, Confocal microscopy image of untransfected ARPE-19 cells stained with antibodies to FLAG (red) and VLT protein (pVLT; green), as a negative control. **b**, Confocal microscopy images of uninfected ARPE-19 cells stained for proteins encoded by VZV gene VLT (pVLT; green) and ORF62 (IE62; red) (left panel), or pVLT (green) and VZV glycoprotein E (gE; red) (right panel) as a negative control. **c**, Confocal microscopy images of VZV pOka-infected and uninfected ARPE-19 cells stained for pVLT (green) and VZV IE62 (red) (left panel) or pVLT (green) and VZV gE (red) (right panel). Nuclei in **a-c** were stained with Hoechst 33342 (blue). Magnification: (**a**) 1000X and 2X digital zoom, (**b**) 1000X and (**c**) 200X. Bars=5 μm (**a**), 10 μm (**b**) and 60 μm (**c**).

**Supplementary Figure 13.**
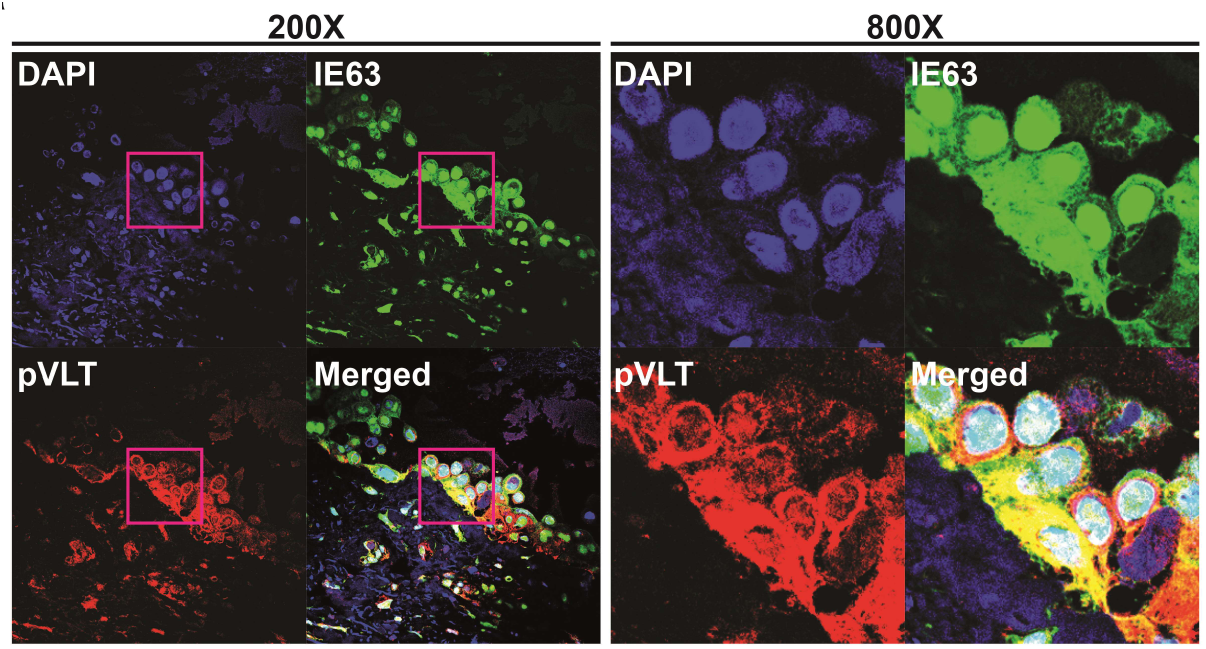
Co-localization of VZV ORF63 and VLT protein in human herpes zoster skin biopsies. Confocal microscopy image showing co-expression of protein encoded by VZV ORF63 (IE63; green) and VLT (pVLT; red) within the same cells of a human herpes zoster vesicle. Nuclei were counterstained with DAPI (blue). Magnification is indicated.

**Supplementary Figure 11.**
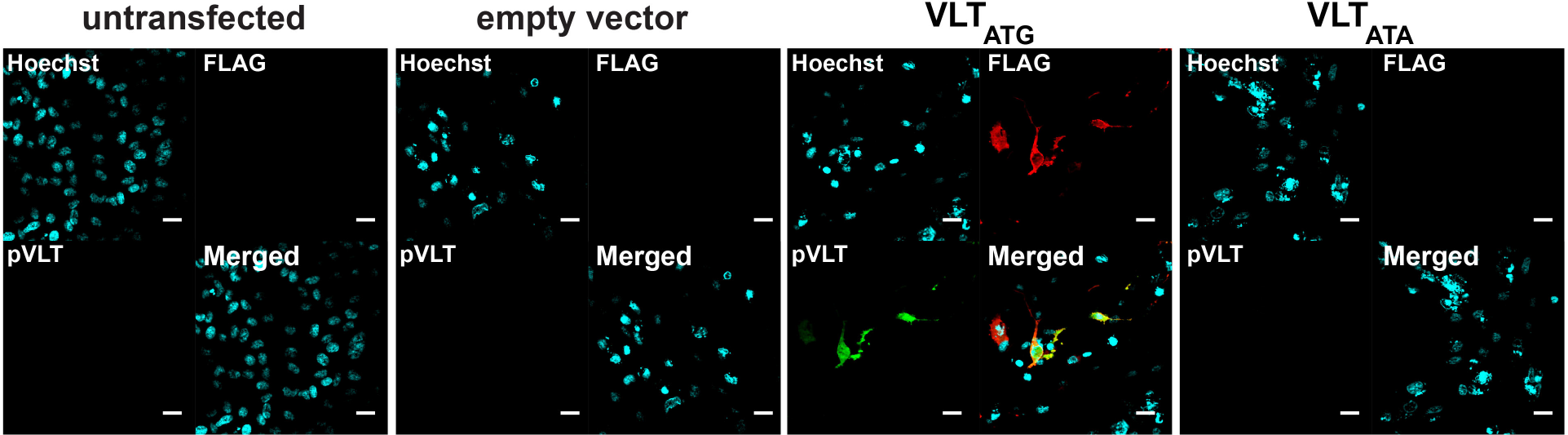
Validation of VLT protein expression in co-transfection experiments. Confocal microscopy image of untransfected ARPE-19 cells and of ARPE-19 cells transiently transfected with plasmids encoding FLAG-tagged VLT (VLT_ATG_), mutated VLT lacking a start codon (VLT_ATA_) or empty control plasmid, in combination with plasmids encoding ORF61, ORF62 and ORF63. Cells were stained using antibodies to FLAG (red) and VLT protein (pVLT; green); nuclei were stained with Hoechst (blue). Magnification: 600X. Bars=20 μm.

## Supplementary Tables

**Supplementary Table 1.** Clinical and virological parameters of human trigeminal ganglion donors used for RNA-sequencing and qPCR. latency transcript.

| Donor<sup>a</sup> | Age (yrs) | Gender | Cause of death | Neurological Disease | PMI<sup>b</sup> (hr:min) | virus gec/10<sup>5</sup> TG cells<sup>c</sup> |  | Virus transcript levels/100 ng RNA<sup>d</sup> |  |  |  |
| --- | --- | --- | --- | --- | --- | --- | --- | --- | --- | --- | --- |
|  |  |  |  |  |  | VZV | HSV-1 | ORF6 3 | VLT2 →3 | VLT3 →4 | LAT |
| 1 | 93 | Fem | Respiratory insufficiency | Alzheimer's Disease | 5:20 | 30 | 30 | 0 | 1.0 | 1.9 | 8606 |
| 2 | 67 | Fem | Heart failure | Alzheimer's Disease | 6:45 | 249 | 26 | 3.2 | 26.8 | 54.7 | 11924 |
| 3 | 81 | Male | Decompensatio cordis | Parkinson's Disease | 5:15 | 164 | 23 | 1.5 | 54.4 | 79.7 | 2078 |
| 4 | 90 | Fem | Sudden cardiac arrest | Alzheimer's Disease | 4:45 | 91 | 7 | 11.9 | 412.4 | 489.0 | 5047 |
| 5 | 67 | Male | Euthanasia | Alzheimer's Disease | 8:00 | 28 | 29 | 0.5 | 4.3 | 9.0 | 8869 |
| 6 | 89 | Male | Sepsis | Alzheimer's Disease | 4:56 | 188 | 0<sup>e</sup> | 8.2 | 131.9 | 188.7 | 1004 |
| 7 | 73 | Fem | Heart failure | Parkinson's Disease | 4:40 | 99 | 13 | 2.4 | 48.4 | 114.7 | 6906 |
| 8 | 83 | Fem | Urosepsis and aspiration pneumonia | Other Neurological Disease | 5:20 | 43 | 553 | 0 | 0.3 | 0.6 | 10739 |
| 9 | 71 | Male | Cardiovascular accident | Parkinson's Disease | 5:05 | 84 | 5 | 0 | 2.3 | 4.8 | 1161 |
| 10 | 71 | Fem | Dehydration and cachexia | Alzheimer's Disease | 4:15 | 66 | 0 | 0.4 | 12.1 | 17.4 | 0 |
| 11 | 69 | Male | Dehydration and cachexia | Alzheimer's Disease | 6:05 | 84 | 119 | 0 | 5.2 | 10.0 | 3094 |
| 12 | 47 | Fem | Euthanasia | Non-demented control | 6:05 | 494 | 0 | 0.3 | 3.6 | 3.3 | 0 |
| 13 | 66 | Male | Uremia/cachexia | Alzheimer's Disease | 7:00 | 26 | 29 | 0 | 0 | 0 | 8205 |
| 14 | 89 | Fem | Euthanasia | Non-demented control | 4:45 | 0<sup>e</sup> | 0 | 0 | 0.9 | 1.2 | 8648 |
| 15 | 80 | Male | Dehydration and cachexia | Other Neurological Disease | 6:35 | 0<sup>e</sup> | 0 | 0.4 | 2.0 | 1.3 | 2699 |
| 16 | 69 | Fem | Euthanasia | Other Neurological Disease | 4:30 | 0 | 42 | 0 | 0 | 0 | 6905 |
| 17 | 95 | Fem | Alzheimer's disease and dehydration | Alzheimer's Disease | 5:30 | 0 | 1 | 0 | 0 | 0 | 24 |
| 18 | 83 | Male | Aspiration pneumonia | Multiple sclerosis | 7:50 | 0 | 10 | 0 | 0 | 0 | 1 |

<sup>a</sup> Donors 1 – 7 were used for RNA-sequencing; donors 1, 3 and 5 were used for miRNA-sequencing; donors 1 – 18 were used for quantitative real-time PCR (qPCR)
<sup>b</sup> PMI: post-mortem interval
<sup>c</sup> Viral DNA load in genome equivalent copies (gec) per 10<sup>5</sup> trigeminal ganglia (TG) cells, as determined by virus-specific and single copy gene hydroxymethylbilane synthase (HMBS)-specific real-time PCR on DNA isolated from about one-fifth part of each TG specimen.
<sup>d</sup> Viral transcript copies per 100 ng total RNA isolated from about four-fifth part of each TG specimen. LAT, HSV-1 latency-associated transcript; ORF63, VZV open reading frame 63; VLT2→3 and VLT3→4, VZV latency transcript (VLT) fragment detected by reversed transcribed real-time PCR using specific primer/probe-combinations.
<sup>e</sup> Individual TG were cut so that approximately one-fifth TG was used for DNA isolation and four-fifth was used for RNA isolation. Occasionally, discrepancies between viral DNA and cDNA qPCRs were observed, most likely due to unequal distribution of HSV-1 among TG neurons and low number of neurons harboring latent VZV. TG positive for either viral DNA or RNA were considered positive for the respective virus, while TG in which no viral nucleic acids could be detected were considered negative for the virus.

**Supplementary Table 2.**
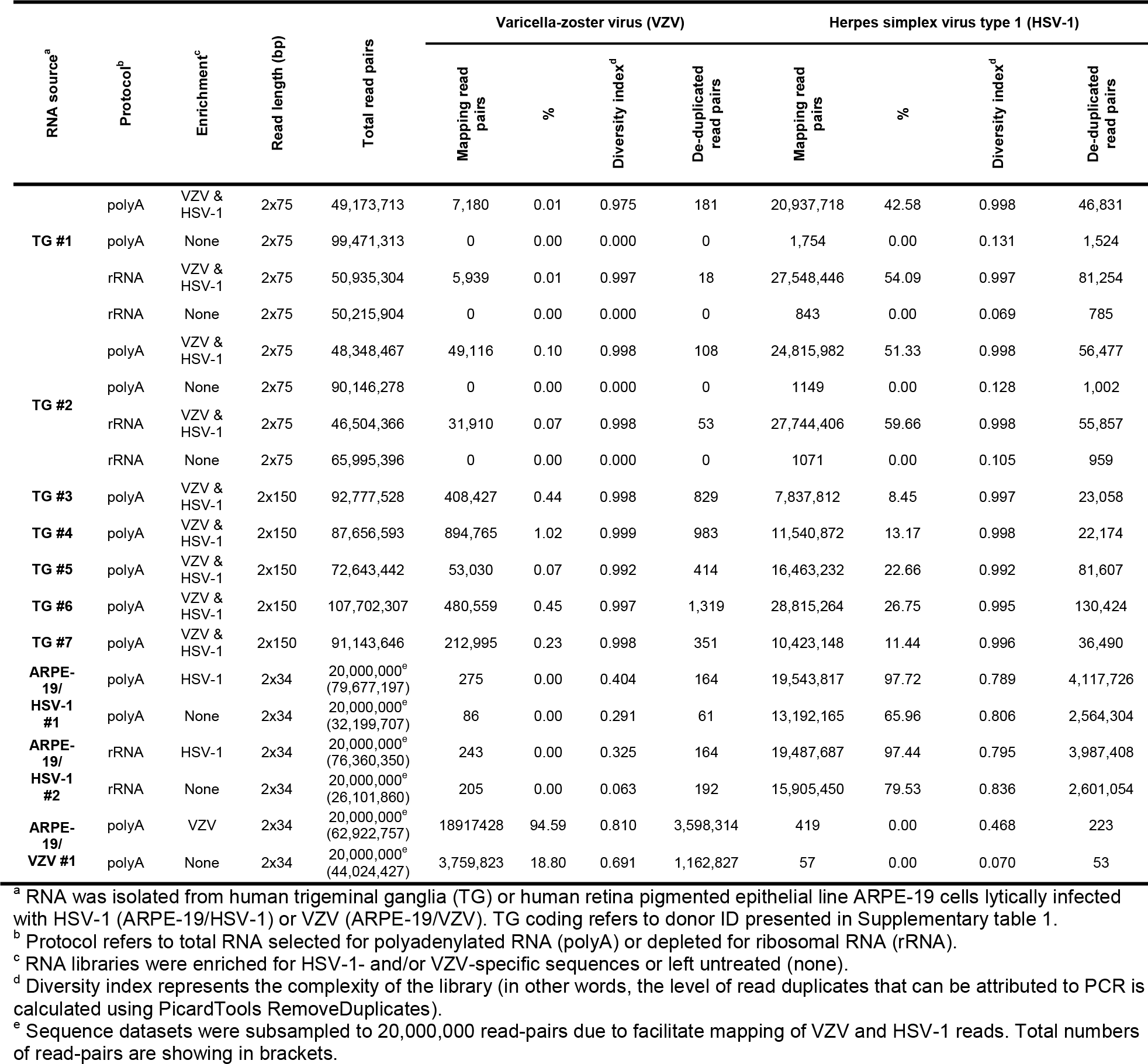
RNA-sequencing metrics.

**Supplementary Table 3.**
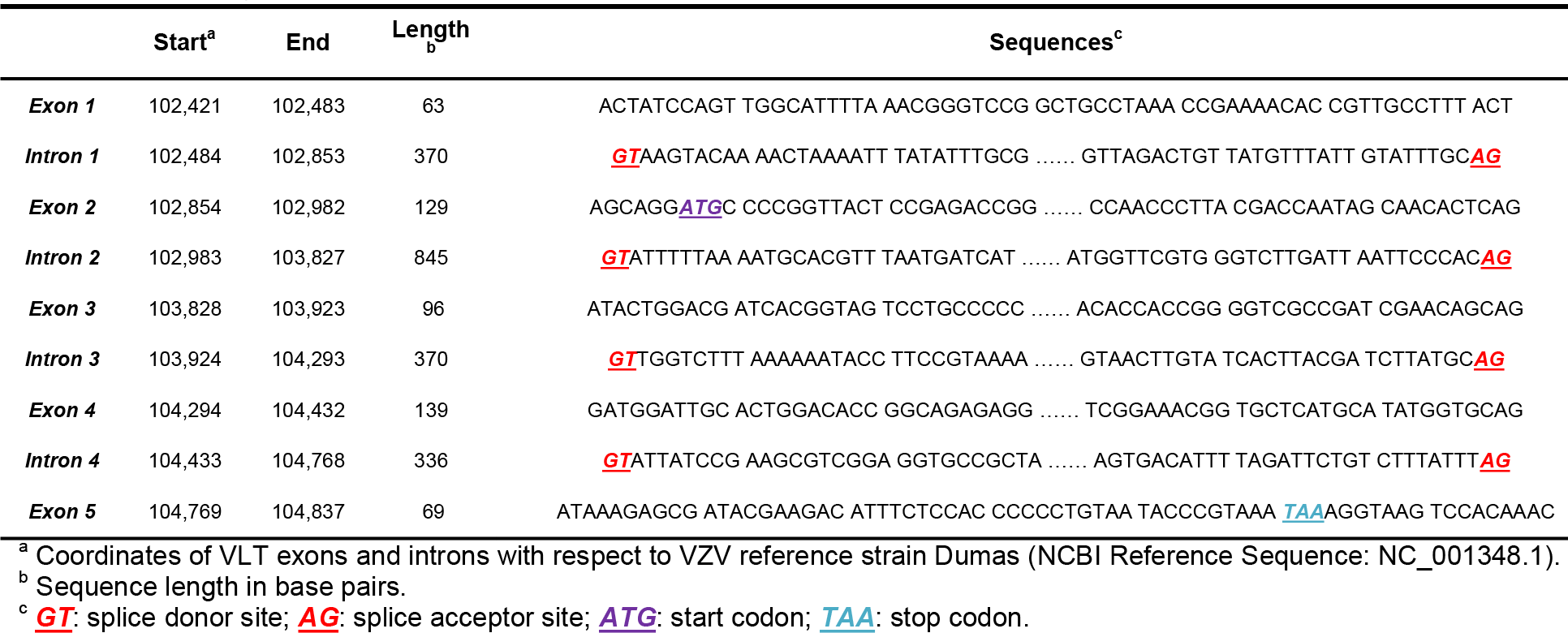
Location and nucleotide sequence of the exons and introns of varicella-zoster virus latency transcript.

**Supplementary Table 4.** Single Nucleotide Polymorphisms (SNPs) across the VLT locus^a^

| Genome position<sup>b</sup> | Reference base<sup>b</sup> | Variant base | ORF | Type of mutation | Codon position | Mutation | Number of genomes<sup>c</sup> | Note |
| --- | --- | --- | --- | --- | --- | --- | --- | --- |
| 102002 | T | C | n.a. | non-coding | . | . | 1 (0) |  |
| 102232 | C | A | n.a. | non-coding | . | . | 1 (0) |  |
| 102268 | T | C | n.a. | non-coding | . | . | 1 (0) |  |
| 102309 | C | A | n.a. | non-coding | . | . | 21 (6) |  |
| 102351 | A | C | n.a. | non-coding | . | . | 19 (6) |  |
| 102401 | G | A | n.a. | non-coding | . | . | 1 (0) | VLT exon 1 |
| 102403 | A | G | n.a. | non-coding | . | . | 1 (0) | VLT exon 1 |
| 102403 | A | C | n.a. | non-coding | . | . | 8 (0) | VLT exon 1 |
| 102458 | A | G | n.a. | non-coding | . | . | 21 (6) | VLT exon 1 |
| 102510 | T | C | n.a. | non-coding | . | . | 2 (0) | VLT intron |
| 102575 | A | G | n.a. | non-coding | . | . | 2 (0) | VLT intron |
| 102596 | C | A | n.a. | non-coding | . | . | 1 (0) | VLT intron |
| 102601 | T | G | n.a. | non-coding | . | . | 19 (6) | VLT intron |
| 102617 | C | T | n.a. | non-coding | . | . | 3 (0) | VLT intron |
| 102684 | A | T | n.a. | non-coding | . | . | 1 (0) | VLT intron |
| 102738 | T | C | n.a. | non-coding | . | . | 2 (0) | VLT intron |
| 102850 | G | A | n.a. | non-coding | . | . | 1 (1) | VLT intron |
| **102908** | **C** | **T** | **VLT** | **non-synonymous** | **1** | **R17T** | **1 (0)** | **VLT exon 2** |
| **102970** | **A** | **G** | **VLT** | **synonymous** | **3** | **Q37** | **2 (0)** | **VLT exon 2** |
| **102977** | **C** | **T** | **VLT** | **non-synonymous** | **1** | **H40T** | **1 (1)** | **VLT exon 2** |
| 103043 | T | C | n.a. | non-coding | . | . | 20 (6) | VLT intron |
| 103169 | A | G | ORF61 | synonymous | 3 | S439 | 1 (2) | VLT intron |
| 103205 | A | G | ORF61 | synonymous | 3 | P427 | 1 (0) | VLT intron |
| 103261 | A | G | ORF61 | non-synonymous | 1 | T409A | 1 (0) | VLT intron |
| 103295 | C | T | ORF61 | synonymous | 3 | G397 | 8 (0) | VLT intron |
| 103322 | T | C | ORF61 | synonymous | 3 | G388 | 1 (0) | VLT intron |
| 103613 | G | T | ORF61 | synonymous | 3 | A291 | 2 (0) | VLT intron |
| 103754 | A | G | ORF61 | synonymous | 3 | G244 | 1 (0) | VLT intron |
| 103778 | A | G | ORF61 | synonymous | 3 | R246 | 2 (0) | VLT intron |
| 103799 | C | T | ORF61 | synonymous | 3 | T229 | 1 (0) | VLT intron |
| 103805 | A | G | ORF61 | synonymous | 3 | P227 | 1 (0) | VLT intron |
| 103955 | A | C | ORF61 | synonymous | 3 | S177 | 1 (0) | VLT intron |
| 104047 | G | A | ORF61 | non-synonymous | 1 | A147T | 1 (0) | VLT intron |
| 104120 | A | G | ORF61 | *synonymous* | 3 | T122 | 1 (0) | VLT intron |
| 104135 | C | T | ORF61 | *synonymous* | 3 | S117 | 1 (0) | VLT intron |
| 104156 | C | T | ORF61 | *synonymous* | 3 | S110 | 1 (0) | VLT intron |
| 104275 | G | A | ORF61 | *non-synonymous* | 1 | D71N | 1 (0) | VLT intron |
| **104403** | **A** | **G** | **VLT** | **non-synonymous** | **2** | **D110G** | **1 (0)** | **VLT exon 4** |
|  | T | C | ORF61 | *synonymous* | 3 | D28 | 1 (0) | VLT intron |
| **104406** | **G** | **A** | **VLT** | **non-synonymous** | **2** | **R111E** | **7 (0)** | **VLT exon 4** |
|  | C | T | ORF61 | *synonymous* | 3 | S27 | 7 (0) | VLT intron |
| 104472 | T | C | ORF61 | *non-synonymous* | 2 | L5S | 1 (0) | VLT intron |
| 104477 | C | T | ORF61 | *synonymous* | 3 | T3 | 1 (0) | VLT intron |
| 104684 | A | C | n.a. | non-coding | . | . | 1 (0) |  |
| 104723 | G | T | n.a. | non-coding | . | . | 1 (1) |  |
| 104737 | T | G | n.a. | non-coding | . | . | 1 (0) |  |
| 104738 | A | C | n.a. | non-coding | . | . | 2 (0) |  |
| 104749 | T | C | n.a. | non-coding | . | . | 1 (0) |  |
| 104865 | G | A | n.a. | non-coding | . | . | 1 (0) |  |
| 104877 | T | G | n.a. | non-coding | . | . | 1 (0) |  |
| 104882 | C | A | n.a. | non-coding | . | . | 11 (0) |  |
| 104898 | A | G | n.a. | non-coding | . | . | 84 (6) |  |
| 104913 | G | A | n.a. | non-coding | . | . | 1 (0) |  |
| 104933 | G | A | n.a. | non-coding | . | . | 4 (0) |  |
| 104939 | C | - | n.a. | non-coding | . | . | 1 (0) |  |

<sup>a</sup> SNPs within the coding region of VLT are shown in bold and within ORF61 are shown using italics.
<sup>b</sup> Reference bases and genome co-ordinates correspond to VZV reference strain Dumas NC\_001348.1
<sup>c</sup> Data originate from a multiple sequence alignment of VZV wildtype (n = 88) and vaccine / vaccine rash (n = 38) strains obtained from Genbank in August 2017. The numbers of wildtype and (vaccine rash) genomes carrying the SNP are reported.

**Supplementary Table 5.**
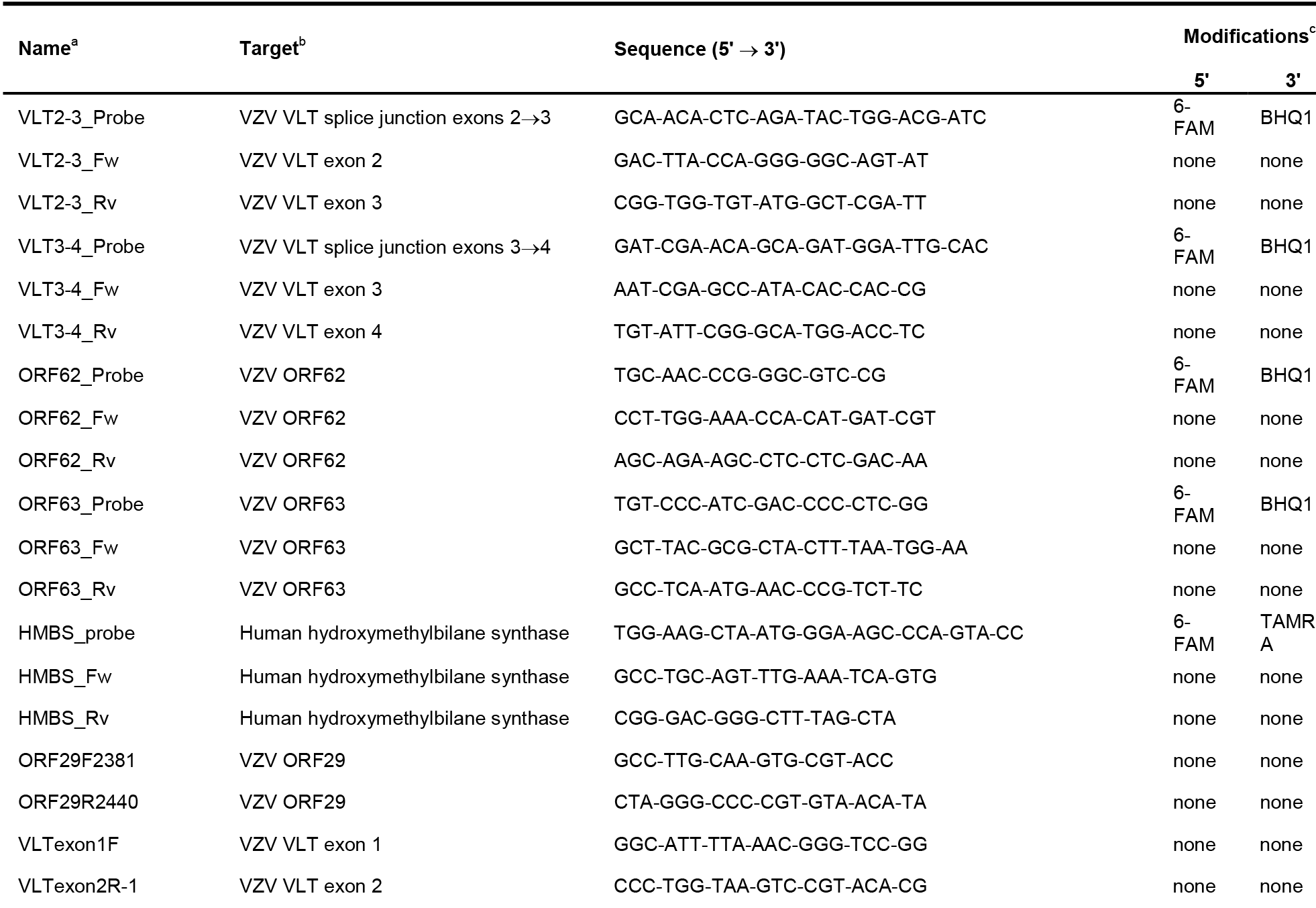
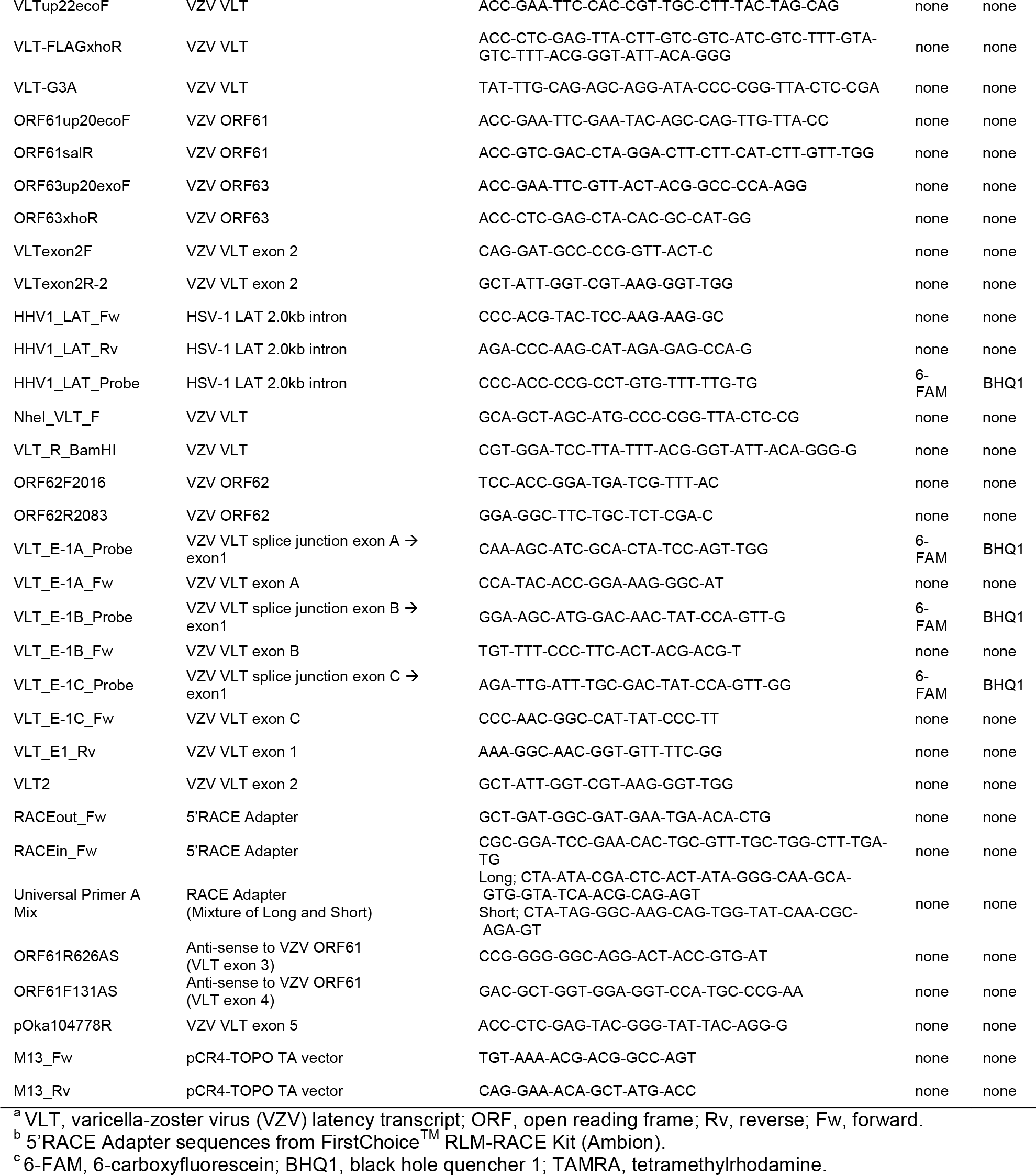
Primers and probes.

## Materials and Methods

### Human clinical specimens

Human trigeminal ganglia (TG), obtained at 6:01 h ± 1:47 hr (average ± SD) after death, were provided by the Netherlands Brain Bank (Amsterdam, the Netherlands) (Supplementary Table 1). All individuals provided prior written informed consent for brain autopsy and the use of material and clinical information for research purposes. All study procedures were performed in compliance with relevant Dutch laws and institutional guidelines, approved by the local ethical committee (VU University Medical Center, Amsterdam, project number 2009/148) and was performed in accordance with the ethical standards of the Declaration of Helsinki. Majority of TG donors had a neurologic disease history affecting the central nervous system (mainly Alzheimer’s and Parkinson’s disease) and cause of death was not related to herpesvirus infections. TG biopsies were either formalin-fixed and paraffin-embedded (FFPE) or snap-frozen in liquid nitrogen and stored at −80°C. Human fetal DRG were obtained from the Academic Medical Centre Amsterdam (the Netherlands) according to relevant Dutch laws and approved by the institutional ethical committee (Erasmus MC, Rotterdam, MEC-2017-009). FFPE punch biopsies of one healthy control subject and five herpes zoster skin lesions were obtained for diagnostic purposes. Zoster biopsies were confirmed VZV DNA positive by virus-specific realtime PCR (qPCR). According to the institutional “Opt-Out” system (Erasmus MC, Rotterdam, the Netherlands), which is defined by the National “Code of Good Conduct” [Dutch: Code Goed Gebruik, May 2011], the surplus human herpes zoster FFPE tissues were available for the current study.

### Cells and viruses

Human retinal pigmented epithelium ARPE-19 cells [American Type Culture Collection (ATCC) CRL-2302] were maintained in 1:1 DMEM (Lonza or Nissui)-Ham’s F12 (Lonza or Sigma-Aldrich) medium supplemented with 8% heat-inactivated fetal bovine serum (FBS; Lonza or Sigma-Aldrich) and 0.6 mg/mL L-sodium glutamate (Lonza or Nacalai). The human MeWo melanoma cell was maintained in DMEM supplemented with 8% FBS and 0.6 mg/mL L-sodium glutamate. Culture of VZV pOka strain and isolation of cell-free virus have been described previously^31^. Low passage clinical isolates VZV EMC-1 and HSV-1 strain F (ATCC VR-733) were cultured on ARPE-19 cells as described^32–34^. All cell cultures and virus infections were performed in a humidified CO_2_ incubator at 37°C.

### Nucleic acid extraction from human TG

Approximately one-fifth of snap-frozen human TG was mechanically dispersed and used for DNA isolation, while four-fifths of the same specimen was used for RNA isolation. DNA extraction was performed using the QIAamp DNA kit (Qiagen) according to the manufacturer’s instructions. For RNA isolation, samples were homogenized in TRIzol (Invitrogen), mixed vigorously with 200 μL chloroform and centrifuged for 15 min at 12,000xg at 4°C. RNA was isolated from the aqueous phase using the RNeasy Mini Kit, including on-column DNase I treatment (Qiagen). DNA and RNA concentration and integrity were analysed using a Qubit Flourometer (ThermoFisher).

### RNA extraction from lytic VZV and HSV-1 infections

ARPE-19 cells were infected with VZV strain EMC-1 by co-cultivation VZV-infected or uninfected cells at a 1:8 cell ratio and harvested in 1 mL TRIzol at 72 hrs post-infection (hpi). Alternatively, semi-confluent ARPE-19 cell layers (75 cm^2^ flask) were infected with HSV-1 strain F at a multiplicity of infection (MOI) of 1 and harvested in 1 mL TRIzol at 24 hpi. RNA isolated as described above was subjected to additional DNase treatment using the TURBO DNA-free kit according to the manufacturer’s instructions (Ambion).

### RNASeq library preparation and sequencing

Four micrograms of total RNA was used as input for the SureSelectXT RNA Target Enrichment protocol (Agilent Technologies G9691 version D0). Here, each sample was either enriched for polyadenylated mRNA (captured by oligo-dT beads, as described by the SureSelect XT protocol) or underwent rRNA-depletion using a NEBNext^®^ rRNA Depletion Kit [Human/Mouse/Rat] (New England Biolabs) according to manufacturer’s instructions. Subsequent to either procedure, captured/remaining RNAs were transcribed to produce cDNA. Fragmentation, cDNA second strand synthesis, end repair, A-tailing and adapter ligation were performed as described in the enrichment protocol. Hybridisation was performed using a modified strategy^20,21^ that incorporated custom-designed SureSelect RNA bait sets for both VZV and HSV-1 in the same reaction at reduced concentration (1:10). Bait sets for VZV and HSV-1 were designed at 12x tiling (i.e. each base position in the genome was covered by 12 distinct 120-mer bait sequences) using custom in-house scripts and are available from the authors upon request. Hybridisation for 24 hrs at 65°C was followed by post-capture washing and optimised PCR-based library indexing (12 cycles for RNA obtained from lytically-infected cultures, 18 cycles for RNA obtained from TGs). Libraries generated from VZV- and HSV-1-infected ARPE-19 cells were multiplexed and sequenced using a 75 cycle V2 high-output kit. Subsequently, three TG were multiplexed sequenced using a 150 cycle V2 mid-output kit followed by a further four TG multiplexed and sequenced using a 300 cycle V2 high-output kit. The decision to use increasing read lengths was informed by the initial discovery of VLT and the desire to better characterise intron splicing.

### Transcriptome mapping and *de novo* transcript reconstruction

Sequence run data were de-multiplexed using bcl2fastq2 v2.17 under stringent conditions (--barcode mismatches 0) and yielded between ∼ 31,000,000 – 107,700,00p0 paired-end reads per sample. Individual sequence datasets were trimmed using TrimGalore software (http://www.bioinformatics.babraham.ac.uk/projects/trim_galore/) to remove low quality 3’ ends and mapped to the human genome (hg19) using BBMap software (http://sourceforge.net/projects/bbmap/) with default parameters. Unmapped read pairs were subsequently aligned against VZV Dumas (NC_001348) and HSV-1 strain 17 (NC_001806) reference genomes deposited at Genbank using BBMap allowing for properly paired reads only to be carried forward in the analysis. Extremely high duplication levels (>0.99) were observed in all enriched libraries generated from TG RNA, an expected feature of enrichment strategies against ultra-low abundance transcripts. Note however that we are not stating that VLT is a low-abundance transcript within a single cell but rather that VZV latently-infected cells are relatively scarce, thus diluting the apparent expression level of VLT. To mitigate this, duplicate reads were removed using Picard Tools MarkDuplicates software (http://broadinstitute.github.io/picard). Resulting assemblies were visualised using a combination of Circos^35^, Artemis^36^, Tablet^37^, SeqMonk (http://www.bioinformatics.babraham.ac.uk/projects/seqmonk/) software and custom R scripts making use of the Rsamtools and Gviz software packages.

To confirm that enrichment for viral nucleic acids did not bias relative levels of viral gene transcription, transcripts per million (TPM) counts were generated using featurecounts (subread package^38,39^) and visualised using scatter plots. High correlation (R^2^ 0.9252 - 0.9678) between enriched and unenriched transcriptomes were observed for both HSV-1 (n=2 biological replicates) and VZV (n=1), as shown in Supplementary Figure S1.

For *de novo* transcript reconstruction, VZV-specific mapping read pairs for each TG were extracted from BAM files and converted to raw fastq format for input into Trinity^40^. Trinity enables transcript reconstruction and, while limited in this case by the scarcity of VZV reads, was able to merge overlapping reads to produce transcript isoforms that, when mapped to the VZV reference genome using BBmap, spanned the four major introns observed in the VZV latency transcript.

### MicroRNA sequencing (miRNASeq) and qPCR profiling

miRNASeq libraries were prepared from 1 μg of total RNA, isolated from human TG and lytically VZV- and HSV-1-infected ARPE-19 cells, using the NEBNext^®^ small RNA Library Prep Set for Illumina^®^ according to manufacturer’s instructions. Libraries underwent 75bp single-end sequencing using an Illumina NextSeq prior to demultiplexing (as outlined above). Sequence reads were adaptor-trimmed using TrimGalore and size selected using BBduk (provided with BBmap) so that only sequence reads between 17-26 bases in length were retained. These were mapped against VZV strain Dumas and the HSV-1 strain 17 reference genomes as well as miRDB using ShortStack (--dicermin 18 --mincov 1 --mismatches 2 --foldsize 200). No putative VZV miRNAs could be detected in either latently infected TGs or lytically infected ARPE-19 cells. By contrast, canonical HSV-1 miRNAs could be detected in both latently infected TGs and lytically infected ARPE-19 cells at high abundance.

### cDNA synthesis and qPCR

cDNA synthesis was performed as described^8^ using 2.0 – 11.7 μg [6.7 μg ± 0.64 (average ± SEM)] of TG-derived total RNA or 5 μg of cell-culture derived DNase-free RNA and Superscript III reverse transcriptase (Invitrogen) with oligo(dT)12-18 primers (Invitrogen). Taqman qPCR was performed in triplicate on DNA and cDNA using Taqman 2X PCR Universal Master Mix (Applied Biosystems) with primer/probe pairs specific for VZV ORF62, ORF63, and the VLT exon 2⟶3, 3⟶4, A⟶1, B⟶1 and C⟶1 junctions and HSV-1 DNA^41^ and LAT, (all from Eurogentec), β-actin (Applied Biosystems) and the human single copy gene hydroxymethylbilane synthase (HMBS) (Supplementary Table 5). Commercially quantified VZVand HSV-1 DNA stocks (Advanced Biotechnologies) and plasmids encoding VZV DNA amplicons (ORF63 and VLT) were used to standardize qPCR reactions and used as positive control in each qPCR. RT-qPCR data were presented as absolute transcript copy number per 100 ng RNA (Fig. 2d and Supplementary figs. 9a and 11a), relative transcript levels defined as 2^−(Ct-value VZV gene − Ct-value β-actin)^ (Figs. 2c and 2e, Supplementary figs. 9b-d), and fold change in gene expression using the 2^−ΔΔCt^ method^42^ in which transcript levels were normalized to β-actin and a reference sample (Figs. 3c and 4a).

## Sequence analysis of lytic VLT isoforms

5’RACE and 3’RACE were performed using the SMARTer^TM^ RACE cDNA Amplification Kit (Clontech) according to the manufacturer’s instructions using total RNA extracted from VZV pOka infected MeWo cells. PCR was performed using KOD-Plus-Ver.2 DNA Polymerase (Toyobo Life Science) templated with 5’RACE ready cDNA using Universal Primer A Mix and ORF61R626AS or pOka104778R, or 3’RACE ready cDNA using Universal Primer A Mix and ORF61R626AS (Supplementary Table 5). Alternatively, 5’Rapid amplification of cDNA ends (RACE) was performed using the FirstChoice^TM^ RLM-RACE Kit (Ambion) according to the manufacturer’s instructions, using 10 μg of total RNA obtained from VZV EMC-1 infected ARPE-19 cells. Nested PCR was performed using Pfu Ultra II Fusion HS DNA Polymerase (Stratagene) and 40 cycles of first round amplification using 5’RACEout_Fw – VLT2-3_Rv primers, followed by 40 cycles of second round amplification using 5’RACEinn_Fw – VLT2_Rv primers (Supplementary Table 5). PCR products were cloned into the pCR^TM^4-TOPO^®^ TA vector (Thermo Fisher Scientific) after addition of 3’A overhang using AmpliTaq Gold DNA polymerase (Thermo Fisher Scientific) or Takara Ex Taq^®^(Takara Bio) and used to transform One Shot^®^ TOP10 competent *E. coli.* Plasmid DNA was extracted using the QIAprep Spin Miniprep Kit (Qiagen), amplified by PCR using M13Fw and M13Rv primers (Supplementary Table 5) and Pfu Ultra II Fusion HS DNA Polymerase (Stratagene). Plasmid DNAs or purified PCR products were sequenced on the ABI Prism 3130 XL Genetic Analyzer using the BigDye v3.1 Cycle Sequencing Kit (both Applied Biosciences) and M13Fw and M13Rv primers. Resultant FASTA sequences were aligned the VZV reference genome Dumas using BBmap, as outlined above. Resultant assemblies were inspected alongside RNASeq data using IGV^43^.

### *In situ* hybridisation

To select for VZV and HSV-1 DNA positive TG tissue areas, DNA was isolated from three consecutive 5 μm FFPE tissue sections using the QIAamp DNA FFPE Tissue Kit and analysed by the respective virus-specific qPCR assays. Subsequently, viral DNA^POS^ FFPE TG specimens, human fetal DRG (negative control) and human zoster skin biopsies were analysed by *in situ* hybridization (ISH) using the RNAScope 2.0 red kit (Advanced Cell Diagnostics) according to the manufacturer’s instructions. In brief, deparaffinised 5 μm tissue sections were incubated with probes designed to cover VZV ORF63 and VLT exons 2 to 5. The probes for the human transcript POLR2A and ubiquitin C (UBC) were used as positive controls and probes specific for the bacterial transcript DAPB were used as negative controls. All probes were designed and produced by Advanced Cell Diagnostics. FastRed was used as substrate to visualize the ISH signal and stained slides were counterstained with haematoxylin and mounted in Ecomount (Biocare Medical). In some experiments, TG sections were incubated with DNase I (Qiagen) or RNAse [Ribonuclease A (25μg/mL) + T1 (25 units/mL; both Thermo Fisher Scientific) diluted in 1xTBS-t], after pre-treatment step #3 for 1 hr at 40°C. To determine the ratio of VZV and HSV-1 transcript expressing TG neurons, slides were scanned using the Nanozoomer 2.0 HT (Hamamatsu) and scored in Adobe Photoshop CS6 (Adobe). Twelve TG from distinct donors were analysed for VZV ORF63 and VLT and 10 TG for HSV-1 LAT, with on average 664 neurons/section (range: 420-1561) and 1-2 sections per TG. Two herpes zoster skin biopsies from distinct donors were analysed with 2-3 sections per donor for each staining.

### Determination of kinetic class of VZV transcripts

ARPE-19 cells (2×10^5^ cells) were infected with pOka VZV cell-free virus (10^3^ plaque forming units) with and without phosphonoformic acid (PFA; 200 µg/mL) for 24 hrs at 37°C. Total RNA was isolated from cells using NucleoSpin RNA in combination with the NucleoSpin RNA/DNA buffer set (Macherey-Nagel) according to the manufacturer’s instructions. Binding DNA was first eliminated from the column in 100 μL DNA elution buffer, the column was treated with recombinant DNase I (5 units/100 μL; Roche Diagnostics) for 1 hr at 37°C and finally RNA was eluted in 60 μL nuclease free water. cDNA was synthesized with 10.4 μL of total RNA and anchored oligo(dT)_18_ primer in a 20 μL reaction using the Transcriptor High Fidelity cDNA synthesis kit (Roche Diagnostics). qPCR following a dissociation curve analysis was performed as described previously ^18^ using SYBR Select Master Mix in a StepOnePlus Real-time PCR system (Thermo Fisher Scientific). The primer sets for β-actin, and VZV ORF61 and ORF49 genes were described previously^18^ and primers for VZV ORF29 (ORF29F2381 and ORF29R2440) and VLT (VLTexon1F and VLTexon2R) are presented in Supplementary Table 5.

### Plasmid construction

The VLT coding sequence (102,468 - 104,818, excluding introns indicated in Supplementary Table 3), ORF61 and ORF63 coding sequences were amplified by PCR of cDNA prepared from pOka-infected MeWo cells showing >80% cytopathic effect using the primer sets VLTup22ecoF and VLT-FLAGxhoR, ORF61up20ecoF and ORF61SalR, or ORF63up20ecoF and ORF63xhoR, respectively (Supplementary Table 5). Products were digested with EcoRI and XhoI (VLT and ORF63), or EcoRI and SalI (ORF61), restriction enzymes and subsequently cloned into pCAGGS-MCS-puro (CAG.Empty) via EcoRI and XhoI sites. The resulting VLT, ORF61 and ORF63 expressing plasmids were named as follows: CAG.VLT-FLAG, CAG.ORF61 and CAG.ORF63. The CAG.VLT(ATA)-FLAG plasmid, in which the ATG start codon of VLT ORF was mutated to ATA to prevent pVLT expression, was generated using primer VLT-G3A with a QuickChange Lightning Multi Site-Directed Mutagenesis kit (Agilent Technologies) according to the manufacturer’s recommendation based on the CAG.VLT-FLAG. All primers used to construct VZV gene expression plasmids are listed in Supplementary Table 5. The pCAGGS plasmid (14) was generous gift from Dr. Jun-ichi Miyazaki (Osaka University, Japan). The pcDNA.ORF62 was a generous gift from Dr. Yasuyuki Gomi (Research Foundation for Microbial Diseases, Osaka University).

### Generation of rabbit anti-pVLT and -IE63 antibodies

Anti-VLT protein (pVLT) antibody was generated by Sigma-Aldrich by immunizing a rabbit with a synthetic peptide encoding the first 19 amino acids of pVLT (MPRLLRDRIAGIPNRVRTY - Fig. S3). The antibody was purified using pVLT peptide conjugated NHS (N-hydroxysuccinimide)-activated sepharose (GE Healthcare Life Sciences). Anti-IE63 antibody was also generated by Sigma-Aldrich by immunizing a rabbit with GST-IE63 protein as described for anti-IE61 antibody^44^. Briefly, GST-IE63 protein was expressed in and purified from *E. coli* BL21 transformed with pGEX-IE63, in which the entire DNA fragment except the first ATG (i.e. ORF63 nucleotide positions 4 - 837) was cloned into pGEX6P-1 bacterial expression vector (GE Healthcare). The anti-IE63 antibody was purified using GST-conjugated NHS-activated sepharose for depleting anti-GST antibody and GST-IE63-conjugated NHS-activated sepharose.

### Immunofluorescent staining and confocal microscopy of cells

The following primary mouse monoclonal antibodies directed to the indicated proteins were used: VZV IE62 (clone 2-B; generous gift from Dr. Yasuyuki Gomi of Research Foundation for Microbial Diseases, Osaka University) and VZV glycoprotein E (clone 9)^45,46^ and anti-DYK (FLAG tag; WAKO). Alexa Fluor 488- and Alexa Fluor 594-conjugated donkey-anti-rabbit and - anti-mouse IgG (Thermo Fisher Scientific) were used as secondary polyclonal antibodies, respectively. Hoechst 33342 (Sigma-Aldrich) was used for nuclear staining. Confocal microscopic analysis were performed as previously described^44^.

### Immunohistochemistry and immunofluorescence on human zoster skin biopsies

Deparaffinised and rehydrated 5 μm FFPE sections of human herpes zoster skin lesions and healthy control skin were subjected to heat-induced antigen retrieval with citrate buffer (pH=6.0), blocked and incubated with mouse anti-VZV IE63 (kindly provided by Dr. Sadzot-Delvaux; Liege, Belgium)^47^, rabbit anti-pVLT or isotype control antibodies overnight at 4°C. Sections were subsequently incubated with biotinylated secondary goat-anti-rabbit Ig or goat-anti-mouse Ig and streptavidin-conjugated horseradish peroxidase (all from Dako) for 1 hr at room temperature. Signal was visualised using 3-amino-9-ethylcarbazole and counterstained with haematoxylin (Sigma-Aldrich). For immunofluorescent staining, Alexa Fluor 488- and Alexa Fluor 594-conjugated goat-anti-mouse and goat-anti-rabbit antibodies were used and sections were mounted with Prolong Diamond antifade mounting medium with DAPI. Confocal microscopic analysis was performed as described^48^.

### Plasmid co-transfection, comparison of viral gene expression by RT-qPCR and protein expression by immunoblotting

Plasmid co-transfection was performed using PEImax (Mw 40,000) (Polysciences, Inc.). High potency linear polyethyleneimine was dissolved in water (1 mg/mL), adjusted to pH=7 with NaOH, filtered through an 0.22 μm filter and stored in aliquots at −20°C until use. CAG.Empty, CAG.VLT-FLAG, or CAG.VLT(ATA)-FLAG (2 μg) with CAG.ORF61, pcDNA.ORF62 and CAG.ORF63 (1 μg) were diluted in knockout DMEM/F12 media (Thermo Fisher Scientific) (50 μL) and PEImax solution (6 μL) were diluted in knockout DMEM/F12 media (50 μL), then both diluent were mixed, left at room temperature for 10 to 15 min to form polyplexes and transfected into ARPE-19 cell (1x10^5^ cells/well in an 12-well plate). Culture medium was changed at 16 hrs post-transfection and cultured for another 48 hrs. Cells were harvested and aliquoted into two fractions. Total RNA extraction from one fraction of transfectants, cDNA synthesis and relative qPCR were performed as described in “Kinetic class of VZV transcripts” section using FavorPrep Blood/Cultured Cell Total RNA mini kit (FAVORGEN BIOTECH) instead of NucleoSpin RNA. The primer sets for VZV ORF62 (ORF62F2016 and ORF62R2083) and VLT (VLTexon2F and VLTexon2R-2) are presented in Supplementary Table 5. Immunoblotting was performed using a rabbit polyclonal antibody against VZV IE61 or IE63, and mouse monoclonal antibodies against alpha-tubulin (B-5-1-2; Sigma-Aldrich), VZV IE62 or FLAG tag (WAKO) for pVLT detection as described previously^44^.

### Data availability

All sequencing runs were performed using an Illumina NextSeq 550 and all demultiplexed fastq dataset are available via the European Nucleotide Archive (http://www.ebi.ac.uk/ena) under study accession PRJEB20723.

## References

1. Kennedy, P. G. E., Rovnak, J., Badani, H. & Cohrs, R. J. A comparison of herpes simplex virus type 1 and varicella-zoster virus latency and reactivation. J. Gen. Virol. 96, 1581–602 (2015).

2. Jones, C. Herpes simplex virus type 1 and bovine herpesvirus 1 latency. Clin. Microbiol. Rev. 16, 79–95 (2003).

3. Nicoll, M. P. et al. The HSV-1 Latency-Associated Transcript Functions to Repress Latent Phase Lytic Gene Expression and Suppress Virus Reactivation from Latently Infected Neurons. PLoS Pathog. 12, e1005539 (2016).

4. Jones, C., da Silva, L. F. & Sinani, D. Regulation of the latency-reactivation cycle by products encoded by the bovine herpesvirus 1 (BHV-1) latency-related gene. J. Neurovirol. 17, 535–45 (2011).

5. Umbach, J. L. et al. MicroRNAs expressed by herpes simplex virus 1 during latent infection regulate viral mRNAs. Nature 454, 780–783 (2008).

6. Cohrs, R. J. & Gilden, D. H. Prevalence and abundance of latently transcribed varicella-zoster virus genes in human ganglia. J. Virol. 81, 2950–2956 (2007).

7. Kennedy, P. G., Grinfeld, E. & Gow, J. W. Latent varicella-zoster virus is located predominantly in neurons in human trigeminal ganglia. Proc. Natl. Acad. Sci. 95, 4658–62 (1998).

8. Ouwendijk, W. J. D. et al. Restricted varicella-zoster virus transcription in human trigeminal ganglia obtained soon after death. J. Virol. 86, 10203–6 (2012).

9. Wang, L., Sommer, M., Rajamani, J. & Arvin, A. M. Regulation of the ORF61 Promoter and ORF61 Functions in Varicella-Zoster Virus Replication and Pathogenesis. J. Virol. 83, 7560–7572 (2009).

10. Moriuchi, H., Moriuchi, M., Straus, S. E. & Cohen, J. I. Varicella-zoster virus (VZV) open reading frame 61 protein transactivates VZV gene promoters and enhances the infectivity of VZV DNA. J. Virol. (1993).

11. Gilden, D., Cohrs, R. J., Mahalingam, R. & Nagel, M. A. Neurological disease produced by varicella zoster virus reactivation without rash. Curr. Top. Microbiol. Immunol. 342, 243–53 (2010).

12. Reichelt, M., Brady, J. & Arvin, A. M. The Replication Cycle of Varicella-Zoster Virus: Analysis of the Kinetics of Viral Protein Expression, Genome Synthesis, and Virion Assembly at the Single-Cell Level. J. Virol. (2009). doi:10.1128/JVI.02137-08

13. Pellet, P. E. & Roizman, B. Chapter 59: Herpesviridae. in Fields Virology (eds. Knipe, D. M. & Howley, P. M.) 2770 – 2802 (Lippincott Williams & Wilkins, 2013).

14. Cohrs, R. J., Gilden, D. H., Kinchington, P. R., Grinfeld, E. & Kennedy, P. G. E. Varicella-zoster virus gene 66 transcription and translation in latently infected human Ganglia. J. Virol. 77, 6660–5 (2003).

15. Nagel, M. A. et al. VZV Transcriptome in Latently Infected Human Ganglia. J. Virol. 85, JVI–01862 (2010).

16. Zerboni, L. et al. Apparent expression of varicella-zoster virus proteins in latency resulting from reactivity of murine and rabbit antibodies with human blood group a determinants in sensory neurons. J. Virol. 86, 578–83 (2012).

17. Haberthur, K. & Messaoudi, I. Animal Models of Varicella Zoster Virus Infection. Pathogens (2013). doi:10.3390/pathogens2020364

18. Sadaoka, T. et al. In vitro system using human neurons demonstrates that varicella-zoster vaccine virus is impaired for reactivation, but not latency. Proc. Natl. Acad. Sci. 201522575 (2016). doi:10.1073/pnas.1522575113

19. Markus, A., Lebenthal-Loinger, I., Yang, I. H., Kinchington, P. R. & Goldstein, R. S. An In Vitro Model of Latency and Reactivation of Varicella Zoster Virus in Human Stem Cell-Derived Neurons. PLoS Pathog. 11, 1–22 (2015).

20. Depledge, D. P. et al. Specific Capture and Whole-Genome Sequencing of Viruses from Clinical Samples. PLoS One 6, (2011).

21. Depledge, D. P. et al. Deep sequencing of viral genomes provides insight into the evolution and pathogenesis of varicella zoster virus and its vaccine in humans. Mol. Biol. Evol. 31, 397–409 (2014).

22. Stevens, J. G., Wagner, E. K., Devi-Rao, G. B., Cook, M. L. & Feldman, L. T. RNA complementary to a herpesvirus alpha gene mRNA is prominent in latently infected neurons. Science (80-.). 235, 1056–1059 (1987).

23. Wang, K., Lau, T. Y., Morales, M., Mont, E. K. & Straus, S. E. Laser-capture microdissection: refining estimates of the quantity and distribution of latent herpes simplex virus 1 and varicella-zoster virus DNA in human trigeminal Ganglia at the singlecell level. J. Virol. (2005). doi:10.1128/JVI.79.22.14079-14087.2005

24. Cheng, Y. C. et al. Mode of action of phosphonoformate as an anti-herpes simplex virus agent. Biochim. Biophys. Acta 652, 90–8 (1981).

25. Jaber, T., Workman, A. & Jones, C. Small Noncoding RNAs Encoded within the Bovine Herpesvirus 1 Latency-Related Gene Can Reduce Steady-State Levels of Infected Cell Protein 0 (bICP0). J. Virol. 84, 6297–6307 (2010).

26. Umbach, J. L., Nagel, M. a, Cohrs, R. J., Gilden, D. H. & Cullen, B. R. Analysis of human alphaherpesvirus microRNA expression in latently infected human trigeminal ganglia. J. Virol. 83, 10677–10683 (2009).

27. Oxman, M. N. et al. A Vaccine to Prevent Herpes Zoster and Postherpetic Neuralgia in Older Adults. N. Engl. J. Med. 352, 2271–2284 (2005).

28. Takahashi, M. et al. Live Vaccine Used To Prevent the Spread of Varicella in Children in Hospital. Lancet 304, 1288–1290 (1974).

29. Cunningham, A. L. et al. Efficacy of the Herpes Zoster Subunit Vaccine in Adults 70 Years of Age or Older. N. Engl. J. Med. (2016). doi:10.1056/NEJMoa1603800

30. Galea, S. A. et al. The Safety Profile of Varicella Vaccine: A 10-Year Review. J. Infect. Dis. 197, S165–S169 (2008).

## Supplementary references

31. Sadaoka, T., Yoshii, H., Imazawa, T., Yamanishi, K. & Mori, Y. Deletion in open reading frame 49 of varicella-zoster virus reduces virus growth in human malignant melanoma cells but not in human embryonic fibroblasts. J. Virol. 81, 12654–12665 (2007).

32. Ouwendijk, W. J. D. et al. Functional characterization of ocular-derived human alphaherpesvirus cross-reactive CD4 T cells. J. Immunol. 192, 3730–9 (2014).

33. Lenac Roviš, T. et al. Comprehensive analysis of varicella-zoster virus proteins using a new monoclonal antibody collection. J. Virol. 87, 6943–54 (2013).

34. Sloutskin, A., Kinchington, P. R. & Goldstein, R. S. Productive vs non-productive infection by cell-free varicella zoster virus of human neurons derived from embryonic stem cells is dependent upon infectious viral dose. Virology 443, 285–293 (2013).

35. Krzywinski, M. et al. Circos: An information aesthetic for comparative genomics. Genome Res. 19, 1639–1645 (2009).

36. Rutherford, K. et al. Artemis: sequence visualization and annotation. Bioinformatics 16, 944–945 (2000).

37. Milne, I. et al. Tablet-next generation sequence assembly visualization. Bioinformatics 26, 401–402 (2009).

38. Liao, Y., Smyth, G. K. & Shi, W. FeatureCounts: An efficient general purpose program for assigning sequence reads to genomic features. Bioinformatics (2014). doi:10.1093/bioinformatics/btt656

39. Liao, Y., Smyth, G. K. & Shi, W. The Subread aligner: Fast, accurate and scalable read mapping by seed-and-vote. Nucleic Acids Res. (2013). doi:10.1093/nar/gkt214

40. Grabherr, M. G. et al. Full-length transcriptome assembly from RNA-Seq data without a reference genome. Nat. Biotechnol. 29, 644–52 (2011).

41. van Velzen, M. et al. Longitudinal study on oral shedding of herpes simplex virus 1 and varicella-zoster virus in individuals infected with HIV. J. Med. Virol. 85, 1669–77 (2013).

42. Livak, K. J. & Schmittgen, T. D. Analysis of relative gene expression data using real-time quantitative PCR and the 2(-Delta Delta C(T)). Method Methods (2001).

43. Robinson, J. T. et al. Integrative genomics viewer. Nat. Biotechnol. (2011). doi:10.1038/nbt. 1754

44. Sadaoka, T. et al. Varicella-Zoster Virus ORF49 Functions in the Efficient Production of Progeny Virus through Its Interaction with Essential Tegument Protein ORF44. J. Virol. 88, 188–201 (2014).

45. Hama, Y. et al. Antibody to varicella-zoster virus immediate-early protein 62 augments allodynia in zoster via brain-derived neurotrophic factor. J. Virol. 84, 1616–24 (2010).

46. Okuno, T., Yamanishi, K., Shiraki, K. & Takahashi, M. Synthesis and Processing of Glycoproteins of Varicella-Zoster Virus (VZV) as Studied with Monoclonal Antibodies to VZV Antigens. Virology 129, 357–368 (1983).

47. Kennedy, P. G., Grinfeld, E., Bontems, S. & Sadzot-Delvaux, C. Varicella-Zoster virus gene expression in latently infected rat dorsal root ganglia. Virology 289, 218–23 (2001).

48. Ouwendijk, W. J. D. et al. T-Cell Tropism of Simian Varicella Virus during Primary Infection. PLoS Pathog. 9, (2013).

